# Development of a universal imaging “phenome” using Shape, Appearance and Motion (SAM) features and the SAM Observation Tool (SPOT)

**DOI:** 10.1101/2025.11.28.691105

**Authors:** Felix Y. Zhou, Adam Norton-Steele, Lewis Marsh, Helen M. Byrne, Heather A. Harrington, Xin Lu

**Affiliations:** Ludwig Institute for Cancer Research, Nuffield Department of Medicine, University of Oxford, Oxford, OX3 7DQ, UK; Lyda Hill Department of Bioinformatics, University of Texas Southwestern Medical Center, Dallas, TX, USA; Novo Nordisk Research Centre Oxford (NNRCO), Innovation Building, Roosevelt Dr, Headington, Oxford, OX3 7FZ, UK; Mathematical Institute, University of Oxford, Woodstock Road, Oxford OX2 6GG, UK; Max Planck Institute of Molecular Cell Biology and Genetics, Dresden, Germany

## Abstract

Cells are plastic, highly heterogeneous and change over time. High-content timelapse imaging promises to reveal dynamic cell behaviors, enabling more accurate identification of cell state and cell fate prediction for biological hypothesis generation and perturbation screens. To empower live-cell imaging based screen, we report the development of 1) a Shape, Appearance, Motion (SAM) “phenome”; a universal set of 2185 image-derived features that act as a image-“transcriptome” to comprehensively quantify an object’s instantaneous phenotype; 2) the SAM-Phenotype-Observation-Tool (SPOT), for image-“sequencing” analysis of phenomes. We validated the effectiveness of unbiased SAM-SPOT workflow on publicly available computer vision and 2D single cell imaging datasets. Importantly, we demonstrated that SAM-phenome outperformed features generated by deep learning AI models trained on >1 million fixed single cell and >5000 single cell video frames, respectively. SAM-phenome and SPOT delivers high-throughput, object-treatment-agnostic, comprehensive screening readouts of dynamics, promising to advance novel molecular target discovery and new medicine development.

## Introduction

Molecular sequencing technologies, particularly scRNA-seq, have transformed our ability to identify previously unknown and rare cell types and states^1–3^. The ability to unbiasedly and simultaneously measure hundreds of transcripts within a cell (average 200–6000 transcripts/cell) regardless of cell and tissue type, to map the transcripts onto a standard reference genome, using widely-accessible tools^4,5^ and implementing a standardized analytical workflow for quantification^6^, enables the systematic identification of known, and the discovery of unknown, genotypes. Unfortunately, scRNA-seq profiles only a single fixed timepoint, and destroys the sample in the process. Therefore, it fundamentally lacks the temporal continuity to assess individual cell trajectories and cell fates, and to filter out the many identified genetic targets that do not impact the final phenotype^7,8^. Moreover, the resolution of scRNAseq analysis is fixed to that of a single cell. Many biological objects, such as multicellular organoids, cannot be measured as a single unit.

In contrast, timelapse imaging, particularly label-free microscopy, is non-destructive and offers unmatched ability to simultaneously capture cell behaviors at both short timescales, with fast acquisition rate, and long timescales, imaging over days and weeks^9^. However, computational analyses of the imaged data lack comprehensive phenotypic readouts that are agnostic of biological process, treatment condition and acquisition parameters. This has restricted studies to either developing simplified analytics targeting a specific or limited set of phenotypes^10,11^ or developing experimental setups that produce binary-like yes/no phenotypic readouts, primarily based on fluorescent tagging of proteins-of-interest^10,12,13^. Unfortunately, the number of simultaneously visualized markers is fundamentally restricted by overlap of excitation and emission spectra and is limited even if spectral unmixing^14^ and temporal encoding^15^ are used. All reporters must further be designed or selected on a per marker basis, and be compatible with live-cell imaging which requires significant specialized knowledge^16–19^. Importantly, all fluorescent reporters photobleach over time, and cells suffer phototoxicity, manifesting artefactual non-physiologically-relevant behaviours^20^. All these underline the unmet need to develop a sufficiently comprehensive direct image-based readout from label-free bright-field live cell time-lapse imaging, suitable for discovering and characterizing diverse phenotypes in a high-content screen. To develop this readout, we took inspiration from scRNAseq.

The success of scRNAseq is due to standardization. First, the prior establishment of reference organism genomes^21–23^ provides a standardized nomenclature, enabling comparison and pooling of scRNA-seq datasets. Crucially, scRNAseq establishes standard experimental protocols to detect, for each cell, the expression of the same standardized ensemble of transcripts comprising the reference transcriptome, without any prior knowledge of the cell type undergoing sequencing^24^. The comprehensiveness of the transcriptome and unbiased detection of all potential transcripts means that scRNAseq is equally applicable to the analysis of known biological function or to the discovery and characterization of novel biological pathways and cellular functions. Moreover, it encourages data reuse, driving the establishment of cell atlases^25–27^. Secondly, the establishment of a standardized simple, modular workflow, comprising clustering, dimensionality reduction, and pseudotime analysis with necessary software tools^4–6^, democratizes scRNA-seq data analysis for non-specialists.

Shape, Appearance and Motion (SAM) are three foundational object characteristics that can be measured directly in video frames to uniquely characterize instantaneous object state. Multidimensional SAM assessment of a moving multicellular object, such as a worm or an organoid, captures image features that can be used to distinguish changes in interactions between cells and those with the surrounding environment^28,29^. A cell’s genetic makeup shapes its ability to respond to environmental stimuli, whilst environmental cues can also sculpt a cell’s signaling networks^8,30,31^. Dynamic cross-talk between environmental perturbations and genetic alterations lead to complex, dynamic, non-linear genotype-phenotype relationships that together determine the ultimate cellular response^32,33^. The ability to measure SAM simultaneously and quantitatively constitute a readily accessible, dynamic, image-based readout that could potentially discriminate phenotypic changes caused by changes in complex genetic circuits or in environmental factors^11,34^. Especially, assessing SAM over long-time horizons would enable more accurate discovery of novel predictors of the fate of a moving multicellular object or organism.

To date, in contrast to scRNA-seq, no standardization exists for measuring and analyzing shape, appearance and motion features of cells. Existing software packages (e.g., CellProfiler^35^, TISmorph^36^) define their own set of phenotypic features, concatenating independently computed image features without systematic design rationale and establishment of effectiveness on limited datasets and phenotypes. The comprehensiveness and general applicability has not been systematically verified. Moreover, being developed primarily for analysing fixed multiplexed images, these software packages provide limited assessment of motion features. Many studies simply compute image features ad-hoc, on a per experiment basis, targeted at hypothesis verification^37–40^. Temporal analysis of computed features presents further unique challenges. Whilst methods are developed to improve identification and tracking of individual cells, there is limited consideration of how to optimally integrate multidimensional SAM features extracted per cell instance with tracking information for the ultimate objective of assessing how perturbation conditions affects the control^17^. Most biological imaging analyses heuristically average the features of tracked objects temporally and/or ensemble-wise within a condition for comparison or resort to training an AI model^40–43^. These practices fail to properly account for inherent heterogeneity of the biological system, including the impact of potential artefacts and temporal changes. Heterogeneity is a hallmark of cells in complex tissue. True biological variation exists between individual cells, and within cells of the same type, in the same microenvironment, and is important for proper function. Indeed, a characteristic signature of diseased tissue is a shift in the observed cell heterogeneity^31,44–46^. Recently, the development of deep learning AI has driven approaches to automatically ‘learn’ the most relevant imaging features directly from input data without explicit rational design^47–50^. There is, however, no large publicly available data repository for SAM method development. Whilst there are now large multiplexed fixed imaging datasets^51,52^, these lack further comprehensive phenotypic annotation to be used like gold-standard computer vision datasets such as ImageNet^53^, or COCO^54^ to benchmark progress and drive systematic AI development. Moreover, due to its specialist nature, we cannot reliably crowdsource annotations as practiced in computer vision^55^. Training any foundational AI to generate universally applicable SAM features would therefore involve a minimum of constructing an exhaustive well-annotated library^56,57^ (e.g., taking into consideration all possible conditions, cell types, treatment conditions and imaging modalities). Meeting this single requirement involves substantial experimental and infrastructure investment^58^, let alone the fact that any trained foundational AI model requires continual retraining to account for changes in experimental set-up^57,59–61^.

To overcome the aforementioned challenges, we take inspiration from scRNA-seq. That is, to harness the power of label-free bright-field live cell time-lapse imaging, we must first develop: 1) a “standard” image-based “phenome” with the same premise as reference genomes: a set of shape, appearance, and motion features that can be universally applied to ‘image-sequence’ the instantaneous state of any object, or cell, acquired by any imaging modality; and 2) a companion standardized analytical framework, SAM phenotype observation tool (SPOT), to universally analyze all computed SAM phenomes in a dataset. The primary aim of the framework being to discover distinct dynamic phenotypic states and the identification of those conditions that cause bona fide phenotypic responses, all whilst taking into account the temporal evolution of phenotypic heterogeneity within individual conditions. Herein, we describe our development of a suitable 2185-D Shape, Appearance and Motion (SAM) phenome and SPOT “phenome-analyzer”. We validate the universality and effectiveness of unbiased SPOT analysis of dynamic imaging phenotypes on small and large publicly available computer vision and 2D live-cell imaging datasets. Importantly, we demonstrate that SAM-SPOT analysis outperformed features generated by training specialist deep learning AI models on both fixed, and live imaging data, in both classifier accuracy and interpretability. Moreover, in a companion paper (*Reviewers are requested to contact the Editor for access)* we also show that SPOT can characterize subtle and complex 3D organoid dynamics from analyzing only 2D projection videos.

## Results

### Constructing a phenome comprising 2185 SAM features to characterize the instantaneous state of a dynamic object (**Figure 1**)

An imaging “phenome” to detect and compare known, unknown, and complex phenotypes in high-content timelapse videos should measure object-agnostic, phenotype-agnostic, image features characterizing an object at multiple spatial scales: global, regional and pixelwise in the current timepoint and its change to the next timepoint. These features should be easy to compute, with minimal assumptions, and discriminate between phenotypic variations in hundreds of thousands of individual moving objects across all timepoints, regardless of the imaged object or the frequency and length of the acquisition (Fig. 1a).

**Figure 1.**
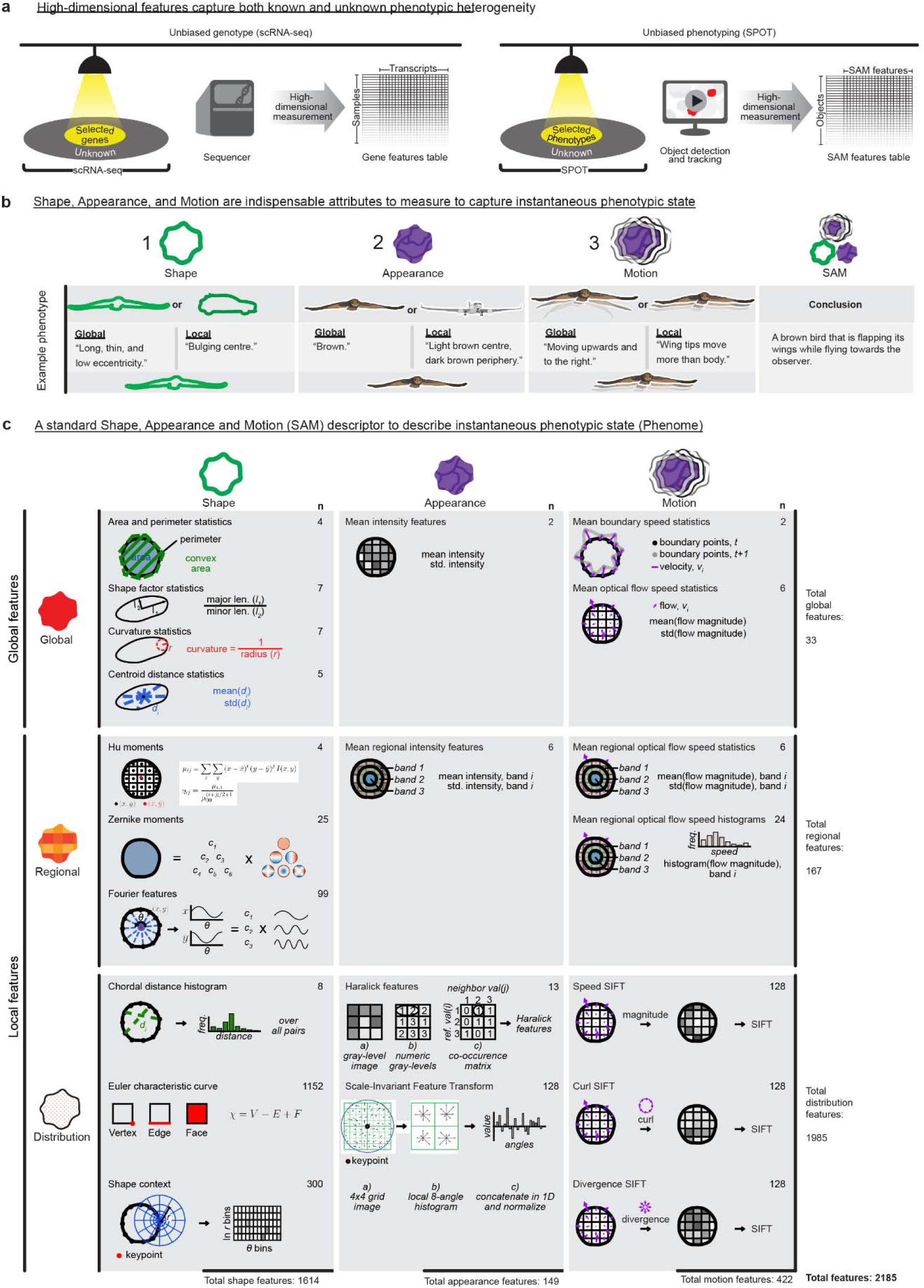
Comprehensive shape, appearance and motion (SAM) characterization enables analysis of phenotypic heterogeneity in timelapse imaging. **a)** Measurement of a standardized high-dimensional library of gene transcripts in single-cell RNA sequencing (scRNA-seq) enables unbiased characterization of the full transcriptome: both that which is known or selected a priori (yellow circle) and unknown (grey ring). Similarly measurement of a standardized high-dimensional set of SAM features could characterize imaging phenotypes that are known or selected a priori (yellow circle) and unknown (grey ring). **b)** Example illustrating how shape, appearance and motion cues provide complementary information and are the minimum three essential concepts necessary to describe the instantaneous state of a dynamic object fully. **c)** Illustration and categorization of the standardized SAM feature set (phenome) used in this study with respect to shape, appearance, or motion (columns) and the spatial scale of object detail they measure (rows). Global refers to the whole object, regional refers to a part of the object, and distribution measures the spatial scale of the object at the level of individual constituent pixels. See Suppl. Table 1 for the definition of individual features. The number of feature dimensions is given assuming an object contour of 200 equidistantly spaced points.

To construct such a phenome, we considered Shape, Appearance Motion (SAM) as constituting a minimal set of image-based object properties to explicitly characterize. We define Shape as an object’s external outline; Appearance as the pattern of image intensity values within the internal object area; and Motion as combined changes in the object’s shape, appearance, and centroid between consecutive timepoints. Whilst there is inevitably feature overlap between the three properties, crucially, they individually provide complementary and indispensable information to fingerprint the object and its instantaneous action state. Fig. 1b depicts how a combination of global and local SAM features enables discrimination between two dynamic objects. Specifically, to ensure a maximally applicable SAM phenome for real data, we established our SAM phenome upon four design principles (Fig. 1c). First, to be comprehensive, SAM properties are measured at multiple spatial scales; global (whole object), local-regional (parts of whole object) and local-(pixel-level) distribution. Second, at each scale, a broad set of SAM features is measured to capture all potential SAM variations, not limited to only identifying object phenotypes within a particular analyzed dataset. This includes measuring redundant features, which characterize the same property but are each computed differently. Redundancy builds robustness to inevitable inaccuracies in segmentation and tracking, whilst still providing additional information. For example, area is significantly more sensitive when an object is partially segmented than the convex area. However, using convex area may underestimate the difference in actual area between two genuinely concave shapes, e.g. a cell with protrusions or that between a convex and concave shape. Third, features are computed to be invariant to object translation or rotation. Evidently, the object is the same under these transformations and therefore should have the same SAM phenome. Reflections of an object are treated as being a different object and therefore have different SAM phenomes. These might reflect differences in the underlying biological process or systematic difference in acquisition (e.g. camera flipped). Last, to enable application to videos of any duration and acquisition frequency, we consider only first order motion features; those computable using a pair of consecutive frames. Higher order motion properties such as acceleration are implicitly captured in tracking over multiple frames.

Briefly, for Shape, we compiled 1614 features, including 23 global (area and perimeter, shape factor, curvature and centroid distance statistics), 131 regional (Fourier features^62,63^, Hu^64^ and Zernike^62,65^ moments) and 1460 distribution features (chordal distance histogram^36^, Euler characteristic curve^66,67^, shape context^68^). For Appearance, we compiled 149 features, including 2 global (mean intensity), 6 regional (regional intensity) and 141 distribution features (Haralick^69^, Scale-Invariant Feature Transform (SIFT)^70,71^). For Motion, we generated 442 features, including 8 global (mean boundary speed, mean optical flow speed statistics), 30 regional (mean regional optical flow speed statistics and histograms) and 384 distribution features (Speed, Curl and Div SIFT). The concatenation of all Shape, Appearance and Motion features (1614+149+442=2185) form our final 2185-D SAM phenome (Fig. 1c and Suppl. Table 1). The rationale for including each feature is provided in Methods. This total number of features is sufficiently high to ensure robustness and discrimination but not so high as to suffer from the curse of dimensionality (see discussion).

To ensure our SAM phenome is computed correctly in code, the following additional processing was also implemented (Methods). Our analytical workflow, SPOT, processes each segmented object as a 200-point polygonal contour, to minimize hard disk memory storage whilst still providing an accurate shape reconstruction (Methods, Suppl. Fig. 1). We rasterize the contour to compute any SAM feature (Fig. 1c, Suppl. Table 1) requiring a binary shape. We further applied principal component analysis to consistently align individual object contours with respect to their major length axis, as additional insurance for computing rotation-invariant shape features.

### Development of SPOT (Figure 2)

To jointly analyze the computed SAM phenomes of all segmented object instances across time from video datasets in an unsupervised, data-relevant and high-throughput, streamlined manner with minimal prior assumptions, we developed the SAM Phenotype Observation Tool (SPOT). The four main stages of the SPOT workflow are summarized in Fig. 2, (details in Methods). **Stage 1** comprises two pre-processing steps: (i) 3D-to-2D projection to convert individual video frames acquired as a 3D z-stack to 2D image frames, and (ii) frame-by-frame registration, to remove any translation motion artefacts such as those caused by microscope stage movements. **Stage 2** comprises (i) detection and segmentation of every object-of-interest at every timepoint, followed by (ii) tracking and filtering to retain only “genuine” detected objects (i.e., those that were tracked consistently for a minimum number of consecutive frames). **Stage 3** is SAM feature computation. For all identified and segmented objects at each timepoint (an object instance), SPOT i) computes the SAM phenome and ii) compiles the SAM phenomes for all objects across all videos in the dataset into a single table for combined analysis. **Stage 4** is the temporal analysis of SAM phenomes, and comprises two parts: part A, the discovery and characterization of phenotype dynamics, and part B, identification of modules of covarying individual SAM features to aid interpretation. For part A: SPOT uses dimensionality reduction similar to scRNA-seq workflows to analyze the compiled SAM phenomes (treating each SAM feature as a genetic transcript equivalent) and to map and quantify the temporal evolution of phenotypic heterogeneity. Specifically, SPOT: i) filters SAM phenomes to remove all noisy and uninformative features, thereby retaining only the subset of SAM features in the phenome most relevant for the experiment. Per feature normalization of retained features by standard scaling then applying power transform to make each feature Gaussian-like. This normalization enables each feature to be treated independently with equal weight. We then perform regularized linear regression with time, to remove features associated with systematic experimental errors and imaging artefacts in the dataset that should not vary with time; ii) constructs the SAM phenomic landscape, applying dimensionality reduction to the filtered and normalized SAM phenomes to generate a reduced 2D coordinate mapping of the spatiotemporal phenomic variation across all object instances (represented by individual points). This landscape colocalizes object instances with similar SAM features from any timepoint and any condition in 2D, which is critical for enabling simplified downstream analysis; namely, iii) constructing SAM temporal phenotype trajectories to compare and cluster conditions based on a population-level summary of temporal evolution of phenotypic diversity, and iv) finding SAM phenotypic clusters to discretely model the landscape for performing detailed characterization of phenotypic heterogeneity dynamics by the v) monitoring of phenotype cluster dynamics: cluster composition over time and the transition probability between clusters. To interpret the SPOT outputs in part A, in part B, SPOT vi) automatically finds SAM modules: groups of individual SAM features that exhibit the same covariation in the dataset, akin to grouping transcripts by biological pathways in RNAseq, (Suppl. Fig. 2a, 2b). Individual SAM modules can be interpreted by their top representative image exemplars in the dataset and the top contributing individual SAM features (Methods). Finally, (vii) the pattern of SAM module expression aids interpretation of phenotype cluster dynamics, (Suppl. Fig. 2b).

**Figure 2.**
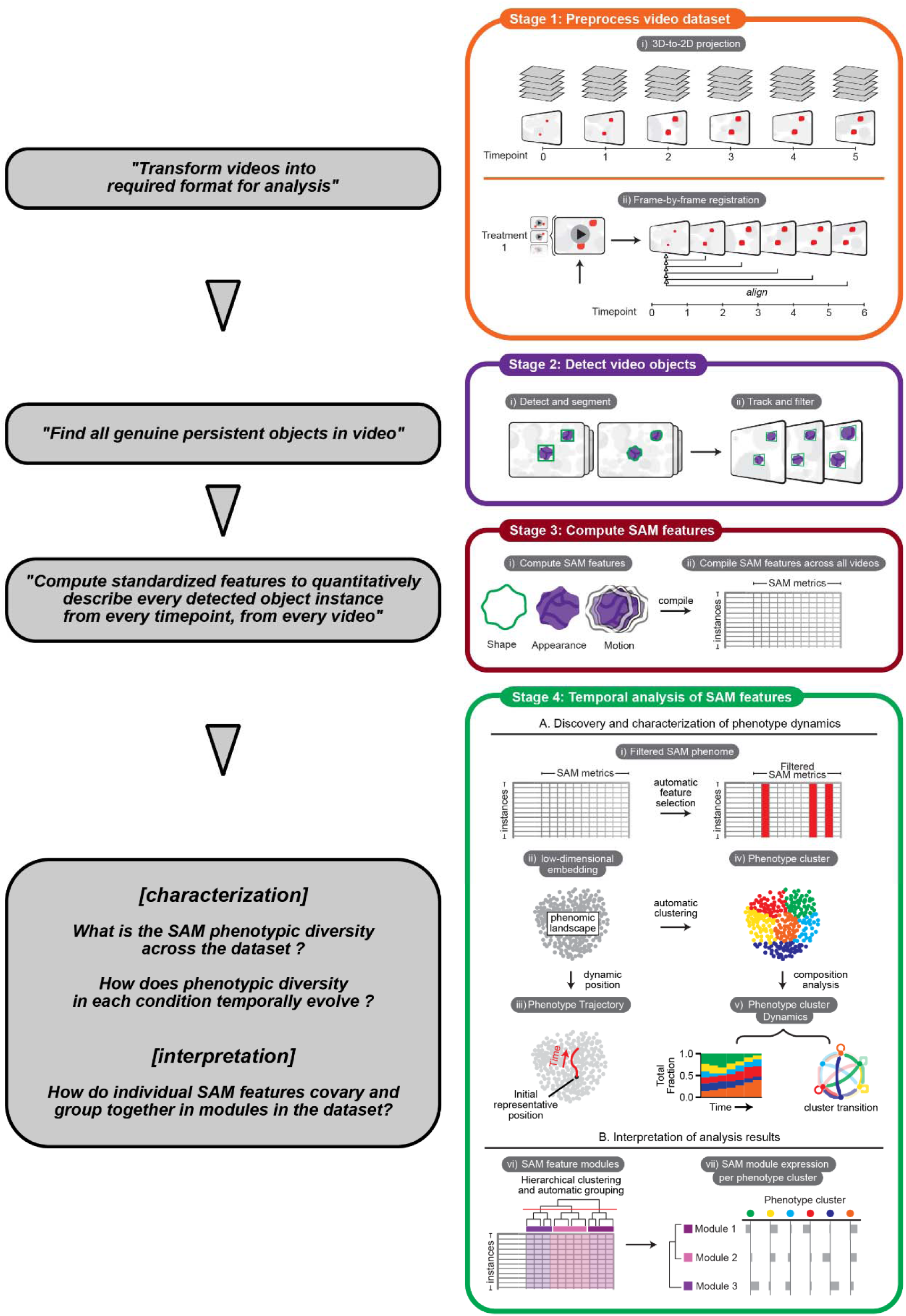
Overview and rationale of the SPOT analysis framework. The four stages in the SPOT workflow with design rationale (left) and illustration of the steps (right) within stage 1: preprocess video datasets (orange box), stage 2: detect video objects, stage 3: compute the SAM phenome for each object instance (brown box) and stage 4: temporal analyses of compiled SAM features (green box).

In summary, the SPOT workflow is designed for high-throughput temporal analysis of objects in videos, enabling the clustering of conditions by phenotypic impact, explicitly accounting for the phenotypic heterogeneity arising from real biological processes and processing artefacts. Treating every temporal instance of an object as an independent observation, SPOT amplifies each object by the number of temporal instances, increasing the ability of analyses to detect under-represented and rare phenotypes. Crucially, this amplification allows SPOT to use dimensionality reduction to map and visualize the full phenotypic variation across space in a 2D coordinate space (the SAM phenomic landscape) in a manner that preserves the global and local SAM similarity between objects, even for small datasets and limited object numbers, for streamlined temporal analysis. Specifically, we use a 2D landscape to construct two simple, universally applicable analytics (population phenotype trajectories and phenotype cluster dynamics) suitable for high-throughput, high-content live-cell imaging screens.

### The SAM phenome comprehensively captures shape and appearance heterogeneity in computer vision datasets with ground truth (Figure 3)

The effectiveness of the constructed SAM phenome and SPOT was tested on two publicly available computer vision datasets with ground truth and benchmarked against CellProfiler^72^ – perhaps the most popular software for measuring shape and appearance features for cell biological applications. To assess the intrinsic information provided by and necessity of all features comprising the phenome, all experiments were conducted without feature selection, applying only standard scaling and a power transform to normalize the features (Methods). Here, SAM phenome refers to the complete ensemble of 2185 SAM features in Fig. 1c and “feature set” to a subset of SAM features in the phenome: shape (SAM-S), appearance (SAM-A) and motion (SAM-M). Dimension (n) refers to the number of SAM features used in an analysis.

**Figure 3.**
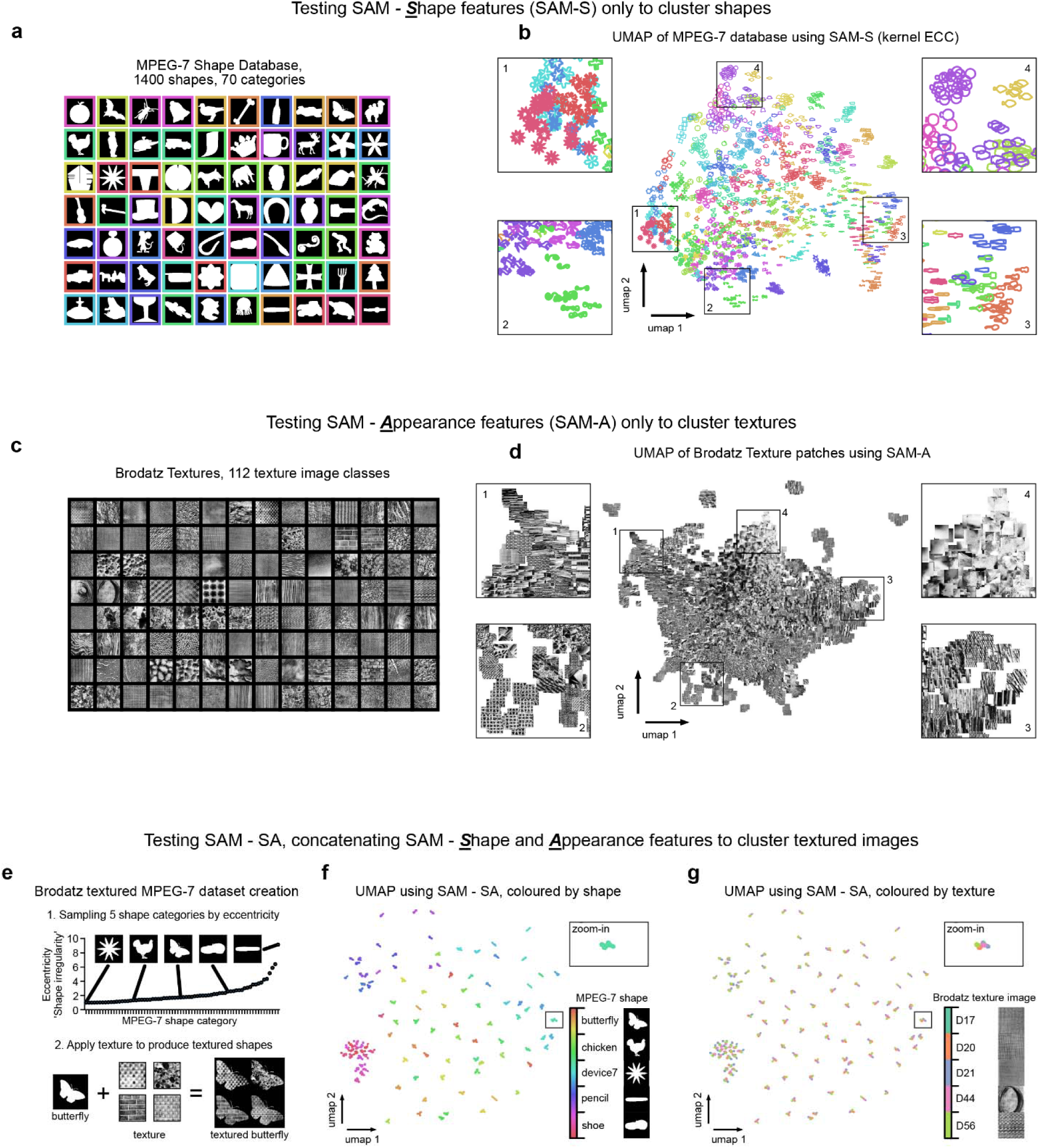
Standardized shape, appearance and motion (SAM) features can comprehensively map phenotypic diversity in computer vision datasets. **a)** Exemplar binary shape images from the MPEG-7 shape computer vision database (one image per shape category depicted), used to test the informativeness of SAM-S, the shape only feature set. **b)** Shape phenomic landscape constructed by applying 2D-UMAP to the full SAM-S (Shape (kernel ECC)) features. Each point is an individual shape image in the MPEG-7 dataset colored by its shape category. Similar shapes colocalize to the same local region of the landscape as shown by zoomed-in images 1-4. **c)** Visual panel summary of the Normalized Brodatz computer vision texture images, used to derive a texture dataset to test the informativeness of SAM-A, the appearance only feature set. **d)** Appearance phenomic landscape constructed by applying 2D-UMAP to the full SAM-A features. Each shape is a 64×64 image patch cropped from the original 512×512 112 Normalized Brodatz texture image. Images cropped from the same original Brodatz texture image and with similar textures colocalize to the same local region of the landscape as shown by zoomed-in images 1-4. **e)** Five representative MPEG-7 shape classes selected according to eccentricity, defined as the ratio of the length of the longest axis to the shortest axis of an ellipse fitted to the shape (**1,** top), used to create textured shapes by combining MPEG-7 binary masks with Normalized Brodatz textures for testing SAM-SA (concatenation of shape and appearance features). The textured shape images were generated by multiplication of images of the same size, one MPEG-7 binary image and one grayscale image crop from a Normalized Brodatz texture image (**2**, bottom). **f)** Shape-appearance phenomic landscape constructed by applying 2D-UMAP to the concatenated full SAM-S (Shape (kernel ECC)) and SAM-A feature sets of the textured shapes created from five selected MPEG-7 shape categories (**e**) and five randomly selected Normalized Brodatz texture images. Each point is a 128×128 image colored by **f)** its source MPEG-7 shape category and **g)** source Brodatz texture.

The MPEG-7 database^73^ is an established computer vision benchmark shape classification dataset containing 1400 shapes equally sampled from 70 different categories (Fig. 3a). Using this dataset, we tested the ability of the SAM-S set to correctly cluster objects into their origin shape categories using unsupervised k-means clustering^74^. Clustering performance was measured based on adjusted mutual information (AMI) and adjusted rand index (ARI) (these scores range between 0 and 1, with 1 being the best performing score) (Methods). We tested the full SAM-S as well as subgroupings of individual SAM-S features (Suppl. Table 2), and applied dimensionality reduction using kernel maps to preprocess Euler Characteristic Curve (ECC) shape features, which has been reported to give better performance^66,75^: SAM-S (kernel ECC) and SAM-S (ECC) indicate the full SAM-S with and without dimensionality reduction, respectively.

2D UMAP analysis applied to SAM-S (kernel ECC) displayed the MPEG-7 shape variation in an ordered manner, from round and symmetrical (left-side of UMAP) to elongated (right-side of UMAP), whilst maintaining clustering amongst individual shape categories (Fig. 3b). 2D UMAP analysis applied to CellProfiler features similarly ordered the MPEG-7 shapes but exhibited a reduced ability to cluster similar-looking shapes, resulting in two distinct point-clouds instead of one (Suppl. Fig 3a,b). Additionally, SAM-S (kernel ECC) (AMI=0.71, ARI=0.52) outperformed subgroupings of SAM-S features and performed better than CellProfiler computed shape features (AMI=0.68, ARI=0.47) (Suppl. Table 2).

The Normalized Brodatz database is an established computer vision benchmark texture (appearance) classification dataset, containing 112 unique texture images^76,77^. Treating these 112 images as independent texture classes, we applied cropping and rotation to construct an augmented dataset of >10,000 image patches. We then tested the ability of the SAM-A set (Fig. 3c, Methods) to correctly cluster patches into their appearance category using unsupervised k-means clustering^74^. As for the shape analysis, the full SAM-A (AMI=0.46, ARI=0.14) outperformed subgroupings of SAM-A features in clustering and was competitive compared with CellProfiler computed appearance features (AMI=0.49, ARI=0.14) (Suppl. Table 2). Notably, despite the similar clustering performance, SAM-A organized image patches with emphasis on textural pattern similarity in 2D UMAP space, whereas CellProfiler emphasized similarities in global brightness (Suppl. Fig 3c,d).

To test the ability to jointly characterize shape and appearance variation, we combined 5 Brodatz textures and 5 MPEG-7 shape classes. For each Brodatz class, we extracted 20 random crops and applied them to each of the 20 unique shape images in each MPEG-7 class, to construct a synthetic dataset of 10,000 images with joint shape and appearance variations (Fig. 3e, Methods). Applying SPOT to compute all combined shape and appearance features (SAM-SA) and using 2D UMAP, we observed that unique shape images were separated as individual tightly grouped colored point-clouds (Fig. 3f). Each point-cloud was further subdivided into 5 smaller point-clouds, each corresponding to a Brodatz texture (Fig. 3f,g and zoom-ins). Thus SAM-SA could reverse-engineer the data generation process. Globally, there is also a broad separation of the 5 primary shape categories, demonstrating SAM-SA finds object shape as the most important global difference. In contrast, UMAP applied to CellProfiler SA-features clusters point-clouds at the coarse level of MPEG-7 shape and Brodatz texture classes but cannot resolve unique shape images (Suppl. Fig. 3e-g). Together these analyses of synthetic datasets confirm that SPOT’s SAM phenome is more discriminative than CellProfiler in capturing joint shape and appearance variations.

### Applying SAM phenome and SPOT to characterize Shape, Appearance and Motion of objects in YouTube videos with ground truth (Figure 4)

We used our SAM phenome to characterize combined variations in shape, appearance and motion of individual objects in the A2D dataset^78^, comprising 3782 YouTube videos of unconstrained ‘real-life’ scenarios. The videos depict seven classes of moving objects (adult, baby, bird, cat, dog, ball and car) performing nine different movements: still (no movement), climbing, crawling, eating, flying, jumping, rolling, running, and walking (Fig. 4a). A single movement class can be performed by different objects, but no object can perform all eight movements, totaling 43 unique action-movement pairings. As expected, SAM phenome significantly outperformed SAM-S, SAM-A and SAM-M features alone in clustering and in training supervised classifiers of object, movement and object-movement pairings (Suppl. Table 2, scores ranging from 0 to 1, with 1 being the highest score).

**Figure 4.**
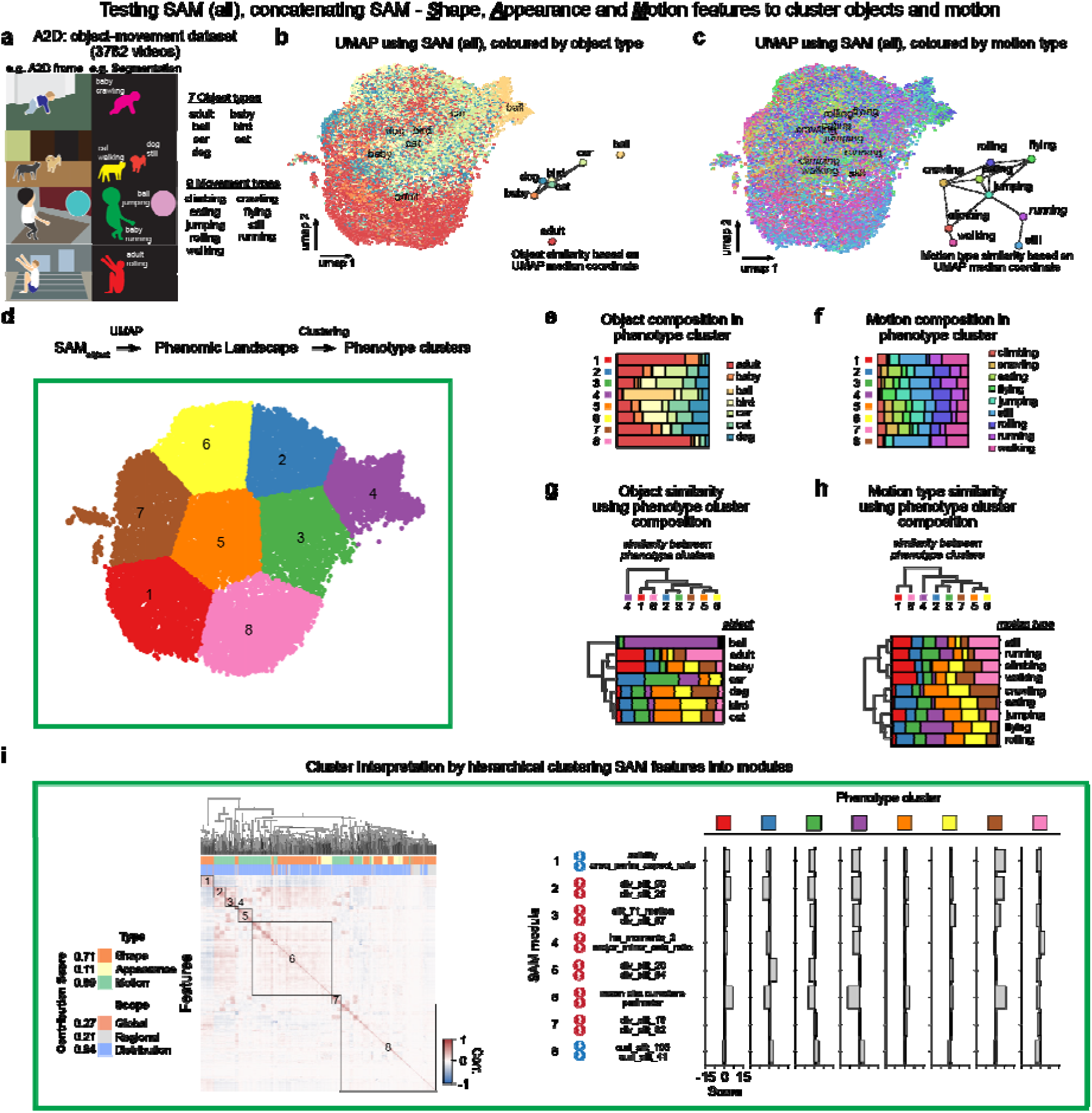
Testing Shape, Appearance and Motion features to cluster objects and motion in the A2D YouTube dataset. **a)** Schematic examples of the object-movement pair annotation in the YouTube-derived A2D dataset used to test the full SAM feature set on videos representing real-life conditions. SAM phenomic landscape of object-motion constructed by applying 2D-UMAP to the full SAM feature set concatenating SAM-S (kernel ECC), SAM-A and SAM-M feature sets colored by **b)** object type or **c)** movement class for each A2D annotated object in the test split. Insets: similarity graph between object and motion classes, respectively, based on the pairwise distance between median 2D-UMAP coordinates. **d)** SAM phenomic landscape constructed by applying UMAP to the SAM phenome, followed by identification of phenotype clusters by k-means clustering of 2D-UMAP coordinates using the elbow method. **e)** Object and **f)** motion composition according to phenotype cluster. **g)** Similarity between object types based on hierarchical clustering of their phenotype cluster composition (bottom). Similarity between phenotype clusters based on hierarchical clustering of their object type composition (top). **h)** Similarity between movement type and phenotype clusters based on hierarchical clustering, same as for g). **i)** Contribution score of shape, appearance and motion, and global, regional and distributional features in explaining the dataset variance defined as the absolute value of the first principal component (Methods) (left). Automated hierarchical clustering of the covariation between SAM features to identify principal SAM feature modules (middle, modules outlined and numbered along the diagonal). The expression of each SAM module (labeled as ‘score’ in barplots, Methods) in each phenotype cluster (right). The top driving SAM features for each module are stated, with arrows indicating whether the feature is enriched (up arrow) or depleted (down arrow).

We next used SPOT analysis to interrogate the phenotypic heterogeneity. As intended by design, SPOT co-clustered object-movement (actor-action) pairings in the UMAP phenomic landscape. The ‘ball’ object maps to the top right of the UMAP, colocalized with the ‘flying’ movement whereas the ‘adult’ object maps to the bottom of the UMAP where it is associated with running, jumping, walking and still – actions most frequently paired with ‘adult’ within this dataset (Fig. 4b,c). Overall, the UMAP spatially arranges object and movement types semantically such that we construct meaningful similarity graphs for object (Fig. 4b) and movement type (Fig. 4c), and joint object-movement pairings (Suppl. Fig. 4a), using only 2D median UMAP coordinates (Methods). This semantic organization was maintained when we varied the choice of dimensionality reduction methods, demonstrating that our constructed SPOT phenome is indeed comprehensive and discriminative (Suppl. Fig. 4b). Herein, we use UMAP in SPOT as the default dimensionality reduction method due to its grounding in mathematical theory^79^, scalable computation, robust open-source code implementation and proven performance in literature. SPOT (Stage 4) partitioned the 2D UMAP landscape into 8 phenotype clusters, with cluster number automatically inferred by the elbow method (Fig. 4d). These clusters should be considered as “prototypical SAM phenotypes”. Consequently, we do not expect each cluster to correspond exclusively to any one single object or movement type (Fig. 4d). However, one object or movement type may be stereotyped, having such distinct phenotypic characteristics that they primarily dominate a cluster or several clusters. For example, adults predominate in clusters 1, 8 (Fig. 4e); and these clusters also have near-identical motion composition patterns of being still or walking (Fig. 4f). Likewise, the predominant object in cluster 4 is ball, followed by car and bird (Fig. 4e), and this cluster expectedly has the largest proportion of flying objects (Fig. 4f). Hierarchical clustering of objects correctly identify ball as most dissimilar to other objects, human adult and baby as a clade, and distinguishes car from animals: dog, bird and cat (Fig. 4g). Movement type is less distinct, due to the diverse shape and appearance of objects conducting the same movement. Nevertheless, the hierarchical clustering is meaningful and finds a grouping of aerial-associated activities: jumping, flying and rolling; vertical activities: walking, climbing, running and still; and a miscellaneous grouping of crawling and eating (Fig. 4h). We can similarly group phenotype clusters by similarity of object and movement type composition (Fig. 4g,h).

To interpret individual phenotype clusters with respect to SAM features, we performed SPOT Stage 4B (Fig. 4i, Methods). Given that SAM phenome annotates each feature by concept (shape, appearance and motion) and by spatial scale (global, regional, distribution), from a grouping of features, SPOT can compute a score of the group contribution to the total phenomic variation (Methods, Suppl. Fig. 2b). In the A2D dataset, we find the primary sources of variation are shape and motion, at (pixelwise) distribution scale with contribution scores of 0.71, 0.69, and 0.94 respectively (Fig. 4i, left). SPOT automatically identifies 8 distinct SAM modules, mutually exclusive groupings of correlated SAM features by hierarchical clustering, and their most relevant image exemplars and individual SAM features. We compute the module expression (the contribution score of each module in each phenotype cluster) to interpret clusters. Module 6, primarily described by mean absolute curvature can be interpreted as measuring the shape ‘complexity’ of an object’s contour. The higher its expression, the more complex the object shape. Accordingly, cluster 4, primarily associated with ‘ball’, a round object, has low module 6 expression. In contrast, cluster 7, comprising a diverse mixture of adult, dog, bird, baby (Fig. 4e) and movement types (Fig. 4f) has the highest module 6 expression. These computer vision results demonstrate that SPOT’s SAM phenome construction is comprehensive and generally applicable to diverse datasets, such that SPOT can automatically find the most relevant subset of shape, appearance and motion features for analysis.

### SPOT comprehensively profiles dynamic phenotypic heterogeneity in single cell videos (Figure 5)

To test the applicability of SPOT workflow (Fig. 2) for biological image analysis, we applied SPOT to detect and compare cell behaviors in live-cell imaging, using two Cell Tracking Challenge^80^ datasets (with reference segmentation and single-cell tracks provided). The first glioblastoma-astrocytoma U373 cell dataset records cell migration without cell division (Fig. 5a, Suppl. Movie 1). Based on provided segmentations and single-cell tracks of U373 cells, SPOT found seven phenotype clusters (Fig. 5b, left, Methods). Consistent with the expected cell morphodynamics in video, cells in cluster 1 (red) and cluster 7 (brown) have a shape composed of a central circular ‘core’, surrounded by individual finger-like protrusions; cells in clusters 2 (blue), 4 (purple) and 5 (orange) have elongated core shapes with asymmetric lamellipodium; and cells in clusters 3 (green) and 6 (yellow) have a single bright, round nucleus distinguished by the absence or presence of lamellipodia. To aid cluster interpretation, we also colored the phenomic landscape using selected SAM features (Fig. 5b, right). The local point density shows that most cell instances map to the centre (Fig. 5b, right panel, first plot), covered by cells with phenotypic features identified in clusters 1, 7 (spherical) and 3, 6 (bright and round nuclei), whilst the left and right extremities (cluster 2, 4, and 5, respectively) comprise eccentric, elongated cell shapes (Fig. 5b, right panel, second plot). The mean intensity and mean speed exhibit no clustering and no cell division events were detected, meaning these are non-discriminative features in this dataset, as expected (Fig. 5b, right panel, third to fifth plots).

**Figure 5.**
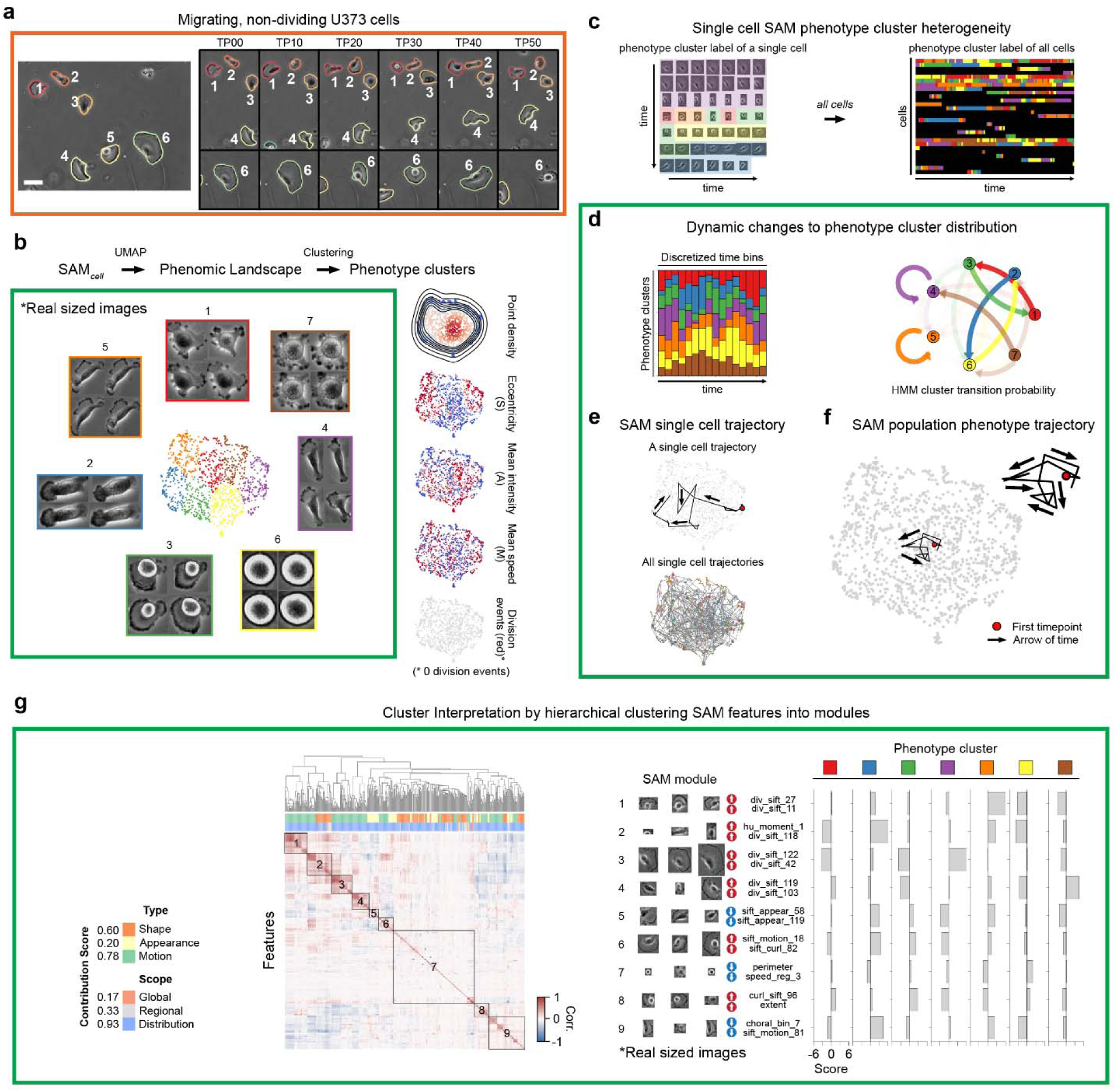
SPOT characterizes phenotypic heterogeneity of single cell migration. **a)** Snapshot of the full field-of-view at timepoint 0 of migrating glioblastoma-astrocytoma U373 cells with the boundary of each cell that is fully in-view in a frame uniquely colored (left), and snapshots of cells at select timepoints (right). The cells did not divide over the video duration. One timepoint is a video frame and is 15 min. Scalebars: 50 µm. **b)** SAM phenomic landscape constructed by applying UMAP to the SAM phenome, followed by identification of phenotype clusters by k-means clustering of 2D-UMAP coordinates using the elbow method. 2×2 image panels show exemplars of the principal phenotypes in each color-coded cluster (left). Local point density of mapped cell instances, whereby each point, representing a cell instance, is colored to indicate low-to-high (blue-to-red) measured values in global SAM features of shape (eccentricity), appearance (intensity), and motion (speed), (right, first to fourth panel, top-to-bottom). Instances were also colored discretely as red or grey to indicate if it was the first timepoint after cell division (right, fifth panel). As the U373 cells did not divide, no point was colored red. **c)** Mapping a single cell tracked over time (left) and all continuously tracked cells (right) into the SAM phenomic landscape of **b)** and coloring each temporal instance by the corresponding phenotype cluster. **d)** Stacked barplot showing the relative frequency of each phenotype cluster over discrete time bins (left). Graph showing the Hidden Markov Model (HMM) inferred transition probability (Methods) of a cell transitioning to another phenotype cluster in the next timepoint, given its phenotype cluster label in the current timepoint (right). Arrows are colored by the source cluster. The more transparent the arrow, the smaller the probability of the transition. **e)** Single SAM phenotype trajectory summarizing the temporal phenotype dynamics of a single U373 cell (top) and the SAM phenotype trajectory of all cells (bottom) with starting timepoint colored red. Black arrow shows the directionality of time. **f)** Single population-level SAM phenotype trajectory summarizing the temporal evolution and phenotypic diversity across all cells. Starting timepoint is colored red. Black arrows show the directionality of time. **g)** Contribution score of shape, appearance and motion, and global, regional and distributional features in explaining the dataset variance defined as the absolute value of the first principal component (left). Automated hierarchical clustering of the covariation between SAM features to identify principal SAM feature modules (middle, modules outlined and numbered along the diagonal). The expression of each SAM module (labeled as ‘score’ in barplots, Methods) in each phenotype cluster (right). Each module is depicted with its top three most representative images, its top driving SAM features and whether the feature is enriched (up arrow) or depleted (down arrow).

The phenotype clusters are SAM analogues of molecular clonotypes^81,82^ from scRNA-seq, decomposing the full phenotypic heterogeneity. To visualize and quantify the temporal evolution of heterogeneity, SPOT assesses the phenotype cluster composition of cell populations within discrete temporal bins (stacked barplots), and infers transition probabilities between clusters by fitting Hidden Markov Models^83,84^ (HMM) to the cluster sequences, constructed by concatenating the phenotype cluster assignment of each temporal cell instance of each tracked single cell (Fig. 5c, Methods). All U373 phenotype clusters expand and contract, most visibly represented by cluster 6 (yellow) (Fig. 5d, left). This periodic behavior is reflective of cluster 6 cells having only a nucleus without lamellipodia transitioning to having large, spreading lamellipodia (cluster 2, blue) and vice-versa (Fig. 5d, right).

To cluster conditions without computing phenotype clusters, suitable for high-throughput, high-content screens, SPOT must derive a single summary ‘signature’ per condition that accounts simultaneously for heterogenous cell behavior and temporal evolution. SPOT does this by adapting pseudotime trajectories^38,85^ from scRNAseq to develop the SAM population phenotype trajectory. For a single tracked cell, we define its phenotype trajectory by chronologically linking the UMAP coordinates of all its individual temporal instances (Fig. 5e, top). The entire set of single-cell phenotype trajectories captures the dynamics of all 29 tracked cells (a total of 1189 cell instances from U373 videos) but is significantly heterogeneous (Fig. 5e, bottom). Crucially, individual trajectories are of different durations, having different start and end times. Generating any consensus trajectory by averaging for any grouping of individual trajectories requires not only complex and time-consuming alignment but may also destroy the heterogeneity pattern. Consequently, SPOT constructs its population trajectory by subdividing time into discrete temporal bins, deriving an average coordinate reflecting the phenotypic diversity of cells in each bin, and linking coordinates chronologically (Methods). The resultant cyclic trajectory of U373 cells: from cluster 1 (red) towards 5 (orange), towards 2 (blue), towards 3 (green), towards 6 (yellow), and back to 1 (Fig. 5f) reflects the periodic expansion and contraction of clusters in stacked barplots and captures the long-time fate of the short-time dynamics revealed by HMM.

The second dataset computer-simulates fluorescently labelled acute promyelocytic leukemia HL60 cells undergoing primarily cell division in-place with negligible cell migration (Suppl. Fig. 5a, Suppl. Movie 1). SPOT generated a SAM phenomic landscape which identified six phenotype clusters (Suppl. Fig. 5b, left), characterizing cells undergoing or having just experienced cell division (clusters 1, 2, 4, 5: high eccentricity reflecting the elongated eccentric shapes during chromosome segregation) and those that have not divided (clusters 3, 6).

As HL60 cells divide, they lose fluorescence. Brighter and darker pixel intensity values therefore discriminate between clusters containing cells that have undergone division more or less recently (Suppl. Fig. 5b,c). Consistent with this, clusters 1 and 3 diminish over time (Suppl. Fig. 5d, left). We also find two short-time HMM transitions representing for cell division at high (cluster 3 to 1) and low (cluster 6 to 2) fluorescence intensity respectively, (Suppl. Fig. 5d, right). Again, individual single cell phenotype trajectories are heterogeneous and it is difficult to extract a consensus behavior pattern (Suppl. Fig. 5e). SPOT’s SAM population phenotype trajectory however clearly reveals the long-time behavior of 4 ‘loop-like’ directional changes (Suppl. Fig. 5f), indicating a total of four cell cycles in the dataset, corroborating the observed division pattern across all individually tracked cells (Suppl. Fig. 5c, right).

SPOT’s SAM module analysis quantifies our qualitative observation that U373 cells primarily migrate and change shape, whereas HL60 cells divide, changing their shape and appearance. For U373 cells, distribution-based shape, motion features were primary contributors of SAM phenotypic variation, with scores of 0.60, 0.78 and 0.93 respectively. (Fig. 5g, left). For HL60 cells, distribution-based shape and appearance features contributed the most, with scores of 0.59, 0.78 respectively (Suppl. Fig. 5g, left). Importantly, appearance (0.20 score) and motion (0.22 score) features were low in U373 and HL60 respectively. Accordingly, SPOT finds more motion-driven modules in U373 (primarily ‘div_sift’, ‘curl_sift’ or ‘sift_motion’ features) (Fig. 5g, right), and more appearance-driven modules in HL60 (‘mean_intensity’, ‘sum_entropy’, ‘diff_variance’, ‘ang_second_moment’, ‘sift_’ features) (Suppl. Fig. 5g, right).

Together, these results confirm that SPOT successfully detects and extracts informative and interpretable SAM features automatically for cellular imaging and can cluster cell instances based on their SAM phenomes. Moreover, using datasets of stereotypical cell migration and division, we validated that SPOT analysis (Fig. 2) generates comprehensive but concise readouts of dynamic phenotypic heterogeneity: capturing the long-time behaviour with stacked barplots of phenotype cluster frequency and SAM population phenotype trajectory; and short-time (i.e. next timepoint) behaviour with HMM inferred cluster transition probabilities.

### SA-phenome outperforms CNN-learnt features on 1.6 million fixed cell images (Figure 6)

To demonstrate that our SAM phenome and SPOT is also discriminative and applicable to large biological datasets, we analyzed two publicly available datasets, one each representing fixed and live-cell imaging. We applied SPOT analysis with SAM phenome and AI neural network learnt features to analyze one of the largest publicly available database of fixed cell imaging, LIVECell (Label-free In Vitro image Examples of Cells)^86^. LIVECell is a well annotated large-scale dataset of over 1.6 million cells imaged using label-free phase contrast microscopy from 8 cell-lines chosen to represent a diverse set of cell morphologies, including small and round, large and flat, neuronal-like with long protrusions, and cultured under different densities^86^ (Suppl. Fig. 6a). The LIVECell dataset is provided as two data splits; a train-validation (val) split (1,011,452 cells) and a test split (415,518 cells) which has cell segmentation masks.

**Figure 6.**
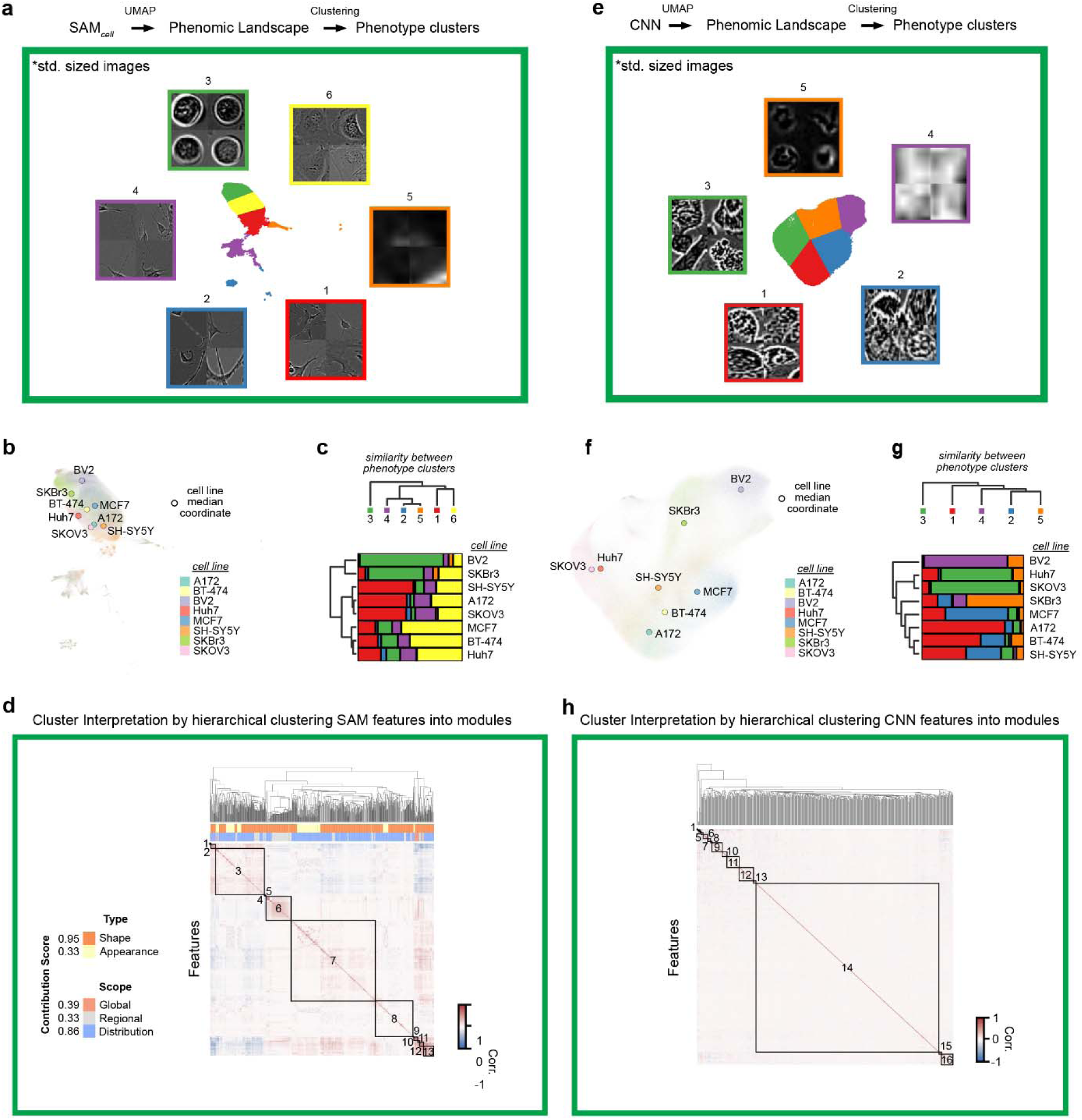
SPOT characterizes phenotypic heterogeneity of different cell types in the LIVECell dataset. **a)** SAM phenomic landscape constructed by applying UMAP to the SAM phenome, followed by identification of phenotype clusters by k-means clustering of 2D-UMAP coordinates using the elbow method. 2×2 image panels show exemplars of the principal phenotypes in each color-coded cluster (Methods). **b)** SAM phenomic landscape with each point (a cell), colored by its cell line. Colored circles show the median coordinate of each cell line. **c)** Hierarchical clustering using average linkage of phenotype cluster based on occurrence across the 8 cell lines (top) and of the 8 cell lines based on their phenotype cluster composition (bottom). **d)** Contribution score of shape, appearance and motion, and global, regional and distributional features in explaining the dataset variance defined as the absolute value of the first principal component. Automated hierarchical clustering of individual SAM features based on pairwise correlation between features to identify SAM modules (middle, modules outlined and numbered along the diagonal). **e)** Phenomic landscape constructed by applying UMAP to CNN features. Cluster coloring and associated exemplar 2×2 image panels as in Fig. 5b. **c)** UMAP with each point (a cell), colored by its cell line. Colored circles show the median coordinate of each cell line**. g)** Hierarchical clustering of phenotype cluster based on occurrence across the 8 cell lines (top) and of cell line using based on their phenotype cluster composition (bottom). **h)** Clustering of individual CNN features into modules based on pairwise correlation between features using hierarchical clustering, as in d).

As LIVECell images are single timepoint snapshots, not videos, we compute the subset of the full SAM phenome comprising only SAM-S shape (kernel ECC) and SAM-A appearance features, (SA-features). SPOT found 709 informative SA-features from the test split dataset (415,518 cells) after feature preprocessing and identified 6 phenotype clusters with three dominant cell appearances: clusters 1,2,4 (neuronal like); cluster 3 (round and nuclear-like); cluster 6 (flat with large lamellipodia) and cluster 5 (image/segmentation artefact) (Fig. 6a). Coloring individual cells in the SPOT UMAP (Fig. 6b), and hierarchical clustering the cell lines using phenotype cluster composition (Fig. 6c) identified 3 main morphological groups: group 1 (small and round shaped [BV2 and SKBr3]); group 2 (cells with distinctive morphology such as elongated neuronal like protrusions [SH-SY5Y & A172]) and large flat lamellipodia [A172 & SKOV3]); and group 3 (cobblestone-like cell clusters [MCF7, BT-474 & Huh7]). This grouping is in complete agreement with the organization of the 8 cell lines in the PCA plot generated using specifically computed morphological features in Fig. 1 of the original LIVECell paper^86^. This highlights the universality of SA-features and the ability of SPOT feature preprocessing to automatically select the most informative subset of SA-features. SPOT identifies 13 SA modules (Fig. 6d, Suppl. Fig. 6c) with shape (0.95) and distribution (0.86) features being the dominant contributors of phenotypic variation.

As an exemplar of AI neural network learnt features, we trained an autoencoder-based convolutional neural network (CNN)^87,88^, which, together with its numerous variants^47–49,89^, represents the most popular method of deriving data-learnt features (Methods). Unlike the rationally engineered SAM phenome, autoencoder-based neural networks, comprise an encoder network which generates a fixed user-specified number of features given an input image, and a decoder networks to reconstruct the input image based on the encoder-generated features. We trained a 512 feature CNN autoencoder using individual image crops of the 1,011,452 cells in the train-val split dataset (Suppl. Fig. 6b). The trained encoder was then applied to the same test-split dataset to generate the 512 CNN features which was analyzed by SPOT identical to using SAM-SA-features. SPOT finds five phenotype clusters, but these identify less diverse and unique cell morphologies than SA-features: cluster 5 (round morphologies); cluster 4 (artefacts), clusters 1,2,3 (all others) (Fig. 6e). Nevertheless, CNN-SPOT’s UMAP does distinguish individual cell lines (Fig. 6f, Suppl. Fig. 7). This suggests that CNN features have likely learnt primarily to distinguish cells by texture and not the finer differences in cell morphology. Consequently, CNN-SPOT’s spatial organization of cell lines in UMAP does not completely agree with the PCA in the LIVECell paper^86^, (Fig. 6f). Critically, hierarchical clustering based on CNN-features finds reduced similarity between the 8 cell lines, grouping only Huh7 and SK-OV-3, and not reflecting what we observe in the images (Suppl. Fig. 6a). Applying SPOT module analysis, CNN-features have high information redundancy. The CNN-SPOT modules lack strong intra-module feature correlation (Fig. 6h) and many have similar representative images (Suppl. Fig. 6d). Critically, unlike the engineered SAM phenome, we cannot ascribe meaning to CNN features (hence they are generically named as Feature 1, 2, 3…etc.), preventing evaluation of the contribution of shape, appearance and motion or further interpretation of individual CNN-SPOT modules. Direct comparison of CNN-SPOT and SA-SPOT on the same dataset illustrates that CNN-SPOT is both less informative and less discriminative than SA-SPOT in identifying subtle morphological similarities between cell lines, notably missing both the neuronal and lamellipodia morphology clusters.

Finally, we also tested the informativeness of SA vs CNN features in training a machine learning classifier to predict cell line. For each, we trained a 2-layer multi-layer perceptron (MLP) on the train-val split and evaluated the MLP performance on the test split, as measured by accuracy and F1 scores (Methods). Both scores range from 0 (no classification) to 1 (perfect classification). SA phenome achieved an accuracy=0.835, F1=0.842, significantly outperforming both the CNN-features without any feature normalization (accuracy=0.544, F1=0.577) used in SPOT analysis, or after feature normalization (accuracy=0.694, F1=0.717) (Suppl. Table 3). The confusion matrix, tabulating the fraction of cells in each cell line (columns) that was correctly predicted (rows), shows that whilst our SA phenome is near-perfect in uniquely identifying cell lines, CNN features confuse a significant fraction of BT-474 as MCF7, and SKOV3 as A172. Based on our SPOT analysis, this likely reflects the inability of the CNN model to learn and generate features capturing finer morphological and textural details.

### SAM-phenome outperforms CNN-LTSM-learnt features on a HeLa cell mitosis video dataset (Figure 7)

To evaluate the full SAM phenome with respect to AI neural network learnt features, we applied SPOT to analyze a well-curated public HeLa cell mitosis video dataset (Fig. 7)^90^. This dataset tracks 326 individual HeLa cells, recording each cell as individual video sequences of 40 timepoints (total 13,040 video timepoints), as its nuclei, visualized by a histone H2B reporter, goes through all 6 distinct stages^91^ of mitosis (Suppl. Fig. 8a). Biologically, mitosis is staged based on the physical state of chromosomes and spindles.

**Figure 7.**
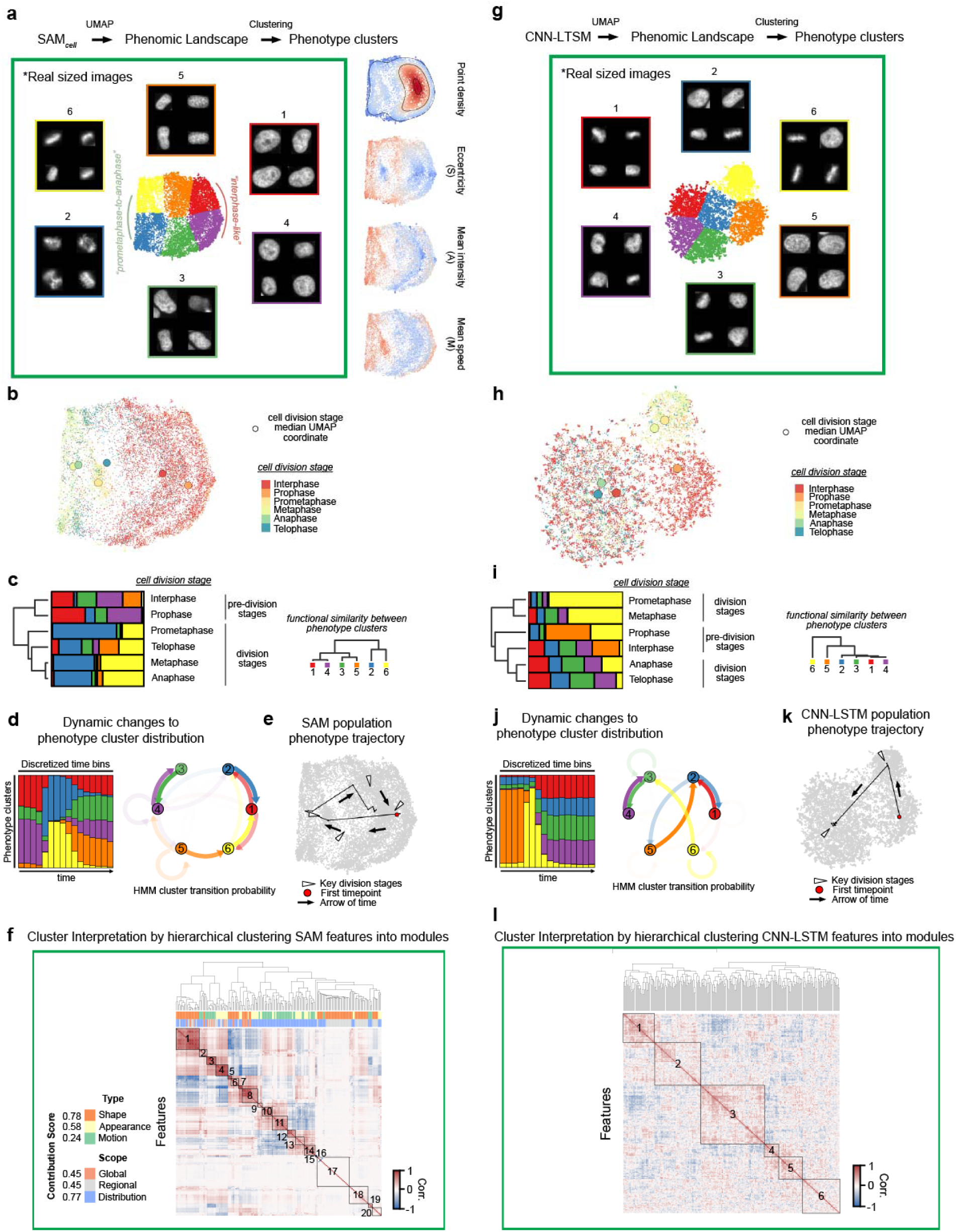
SPOT characterizes nuclei appearance during cell division. **a)** SAM phenomic landscape constructed by applying UMAP to the SAM phenome, followed by identification of phenotype clusters by k-means clustering of 2D-UMAP coordinates using the elbow method. 2×2 image panels show exemplars of the principal phenotypes in each color-coded cluster (left, Methods). Local point density of mapped cell instances, whereby each point, representing a cell instance, is colored to indicate low-to-high (blue-to-red) measured values in global SAM features of shape (eccentricity), appearance (intensity), and motion (speed), (right, first to fourth panel, top-to-bottom). **b)** SAM phenomic landscape with each point (a cell), colored by its cell line. Colored circles show the median coordinate of each cell line. **c).** Hierarchical clustering using average linkage of the 6 stages of cell division based on their SAM-SPOT phenotype cluster composition (left) and of phenotype cluster based on their occurrence across the 6 phases of cell division (right). **d)** Stacked barplot showing the relative frequency of each SAM-SPOT phenotype cluster over discrete time bins (left). Graph showing the Hidden Markov Model (HMM) inferred transition probability of a cell transitioning to another phenotype cluster in the next timepoint, given its phenotype cluster label in the current timepoint (right). Arrows are colored by the source cluster. The more transparent the arrow, the smaller the probability of the transition. **e)** All cell population-level SAM phenotype trajectory summarizing the temporal evolution and phenotypic diversity across all cells and all videos. Starting timepoint is colored red. Black arrows show the directionality of time. **f)** Contribution score of shape, appearance and motion, and global, regional and distributional features in explaining the dataset variance defined as the absolute value of the first principal component. **g)** Phenomic landscape constructed by applying UMAP to the CNN-LSTM features. Cluster coloring and associated exemplar 2×2 image panels as in Fig. 7a. **h)** Phenomic landscape with each point (a cell), colored by cell division stage. Colored circles show the median coordinate of each stage**. i)** Hierarchical clustering of cell line using average linkage based on their CNN-LSTM-SPOT phenotype cluster composition (left), and of phenotype cluster based on occurrence across the 8 cell lines (right). **j)** Stacked barplot of CNN-LSTM-SPOT phenotype cluster composition over time (left), Hidden Markov Model (HMM) transitions (right) as in Fig. 7d. **k)** All cell population-level CNN-LSTM-SPOT phenotype trajectory as in Fig. 7e with starting timepoint colored red and black arrows showing the directionality of time. **l)** Clustering of individual CNN features into modules based on pairwise correlation between features using hierarchical clustering (left).

Here, our aim was to establish whether our universal SAM phenome could detect each stage. Unlike LIVECell, this dataset does not provide ground-truth segmentations. Consequently, we first performed SPOT stage 2, to segment nuclei in every frame by intensity-based thresholding (Methods). To ensure one video sequence corresponds to only one tracked cell, we retained the more centrally located of the two daughter nuclei after cell division. SAM-SPOT identified 186 SAM-features after preprocessing, and found 6 phenotype clusters (Fig. 7a), which appear to capture the changing chromosome appearance during mitosis: clusters 1,4 (interphase-like); clusters 2,6 (prometaphase-to-anaphase); and clusters 3,5 (intermediate-looking). SAM features and the organization of mitosis stage in the UMAP support this pattern (Fig. 7a, right, Fig. 7b). The majority of cell instances are in interphase, characterized by low eccentricity (i.e. circular nucleus) and low mean intensity, and is mapped to the right-side of the UMAP. In metaphase, chromosomes separate to polar ends, leading to elongated shape (high eccentricity), bright (high mean intensity) and large movements (high mean speed), mapping to the left-side of the UMAP. Notably, the median UMAP coordinate of each stage arranges clockwise in a cycle (Fig. 7b) whose separation distance predicts that the greatest SAM characteristic changes occur within the prophase-to-prometaphase and telophase-to-interphase transitions, consistent with observing larger changes in nuclear appearance in video sequences. As validation, hierarchical clustering of mitosis stage based on phenotype cluster composition identified a pre-cell division-related (interphase and prophase) and division-related (prometaphase, telophase, metaphase and anaphase) grouping (Fig. 7c). Moreover, clustering of phenotype clusters by mitosis stage composition generated three pairings of phenotype clusters that functionally partitions of the 2D-UMAP into 3 vertical regions left-to-right, forming a gradient prophase-to-anaphase to interphase-like (Fig. 7b). The SAM-SPOT stacked plot clearly highlights the birth, expansion and disappearance of division-associated clusters 2,6 over time (Fig. 7d, left). The HMM cluster transitions infer that these two clusters originate from cluster 1 and transition back to cluster 1 (Fig. 7d, right). For additional validation, we expect clusters with less defined fate to transition to many clusters, and clusters with predetermined fate to transition to fewer clusters. Accordingly, we observe a lower Shannon entropy of cluster transition for prometaphase-to-anaphase clusters 2, 6 than interphase-like clusters 1, 4. The SAM population phenotype trajectory reveals the entire cycle of mitosis on the 2D UMAP (Fig. 7e), validating the implied clockwise transition from right-to-left side of the UMAP back to right-side of Fig. 7b, and identifies 4 prominent turning points corresponding to key appearance changes at prometaphase; metaphase/anaphase; telophase/interphase and interphase/prophase (Fig 7e). SAM module analysis computes the expected decreasing contribution of shape (0.78), appearance (0.58) and motion (0.28) features to phenotypic variation. Whilst distribution features remain most informative, we found that the contribution scores of global/regional features were highest in this dataset out of all datasets studied. This is due to the evident appearance change from round to elongated nuclei (Suppl. Fig. 8a). SAM-SPOT generated 20 modules with strong intra-feature correlation. Importantly, representative images and features demonstrate that SAM-SPOT modules unbiasedly capture all prototypical nuclei shape-appearances in mitosis (Fig. 7f)

As an exemplar of AI neural network learnt features, we trained an autoencoder-based CNN-LSTM^92–95^ (Convolutional Neural Network-Long Short Term Memory) network to generate features for an input video frame. This network modifies the fixed imaging autoencoder CNN network used for LIVECell dataset for videos, by incorporating an additional bidirectional LSTM layer^95,96^, enabling the CNN-LSTM encoder to generate 256 CNN-LSTM features with a temporal context of 5 frames (Methods, Suppl. Fig. 8b). The number of features was chosen to be comparable to the 186 SAM-SPOT features. As for LIVECell analysis, to ensure like-for-like comparison with SAM-SPOT, the CNN-LSTM was trained using and applied to all 326 video sequences (Methods). CNN-LSTM-SPOT found 6 phenotype clusters, however, compared to SAM-SPOT, individual exemplar images within a cluster showed inconsistent shape-appearance (Fig. 7g). The phenomic landscape indicates that this is due to CNN-LSTM features not capturing the morphological difference of the nucleus being circular in interphase and elongated during prometaphase/metaphase (Fig 7h). Unsurprisingly, clustering of mitosis stage by phenotype cluster composition failed to separate pre-division and division stages (Fig. 7i). The stacked plot does detect clear emergence of elongated nuclei as the cell divides (cluster 6, Fig. 7j, left) and the HMM analysis shows its transition to cluster 3 post-division (Fig. 7j, right), but unlike SAM-SPOT, it fails to explain how cluster 6 must have emerged due to transitions from cluster 5 (round nuclei, pre-division). The CNN-LSTM population phenotype trajectory has only two turning points, verifying that CNN-LSTM features are insufficiently informative of mitosis stage, recognizing only the coarse morphology changes between round and elongated nuclei (Fig. 7k). SPOT module analysis also indicates that this is the primary drawback of CNN-LSTM-features, which generated only 6 SAM modules (Fig. 7l, Suppl. Fig. 7f), and captured only a subset of the phenotypic diversity of SAM-SPOT (Fig. 7f, Suppl. Fig. 7e).

Finally, we also assessed the possibility of training a classifier using SAM-phenome and CNN-LSTM features to classify mitosis stage (Methods). As predicted by SPOT analysis, SAM-phenome (accuracy=0.649, F1=0.708) significantly outperformed CNN-LSTM-features without (accuracy=0.460, F1=0.529) and with additional feature normalization (accuracy=0.454, F1=0.509), (Suppl. Fig. 8g,h; Suppl. Table 4). Confusion matrix analysis shows that whilst the SAM-phenome can clearly identify each division stage, CNN-LSTM features significantly confuses stages, notably unable to distinguish metaphase from prometaphase and anaphase. All these demonstrate that despite explicit incorporation of longer-time temporal dependencies with LSTM, SAM-phenome still unexpectedly outperforms CNN-LSTM features by providing both a more comprehensive and accurate characterization of nuclear morphodynamics in terms of shape, appearance and motion and in training a superior mitosis stage classifier.

## Discussion

A major technological barrier which prevents the effective application of label-free timelapse imaging to biological discovery is a lack of standardization with regard to the measurement and temporal analysis of shape, appearance, and motion (SAM) characteristics of imaged objects. To overcome this barrier, we took inspiration from scRNA-seq analysis workflows^6^ has inspired similarly streamlined analysis of spatial transcriptomics^97^, and developed a 2185-feature SAM phenome and companion standardized workflow, SPOT, for “temporal image sequencing analysis”. Importantly, we demonstrated its universality and effectiveness by analyzing diverse public computer vision and biological datasets, including both small and large-scale datasets, notably A2D^78^, a challenging dataset of >350 YouTube videos; the LIVECell^86^ dataset, with >1.5 million label-free image cells; single-cell tracking datasets^98^; and 326 video sequences (>5000 frames) of the nucleus during cell division^90^.

Rational construction of the SAM phenome was critical to this success (Fig. 1, Methods). A sufficient number of image features is necessary to be 1) comprehensive; characterizing the properties of shape, appearance, and motion at multiple spatial scales (global, regional and pixelwise (i.e. distribution)); and 2) robust to artefacts from image acquisition, segmentation and tracking; achieved by computing redundant features with complementary information e.g., computing both area and convex area. However, excessive features make each SAM phenome too ‘unique’, and hamper pattern finding. As such, our final full 2185-feature phenome represents a good balance. This principle can be best seen in our benchmarking of different subgroupings of SAM-S features on MPEG-7 shapes (Suppl. Table 2). The full SAM-S (kernel ECC) phenome (ranked 1^st^, Suppl Table 2) outperformed all subgroupings, but only due to preprocessing of ECC features (n=1152, 11th) to reduce its dimensionality to 100 features before concatenation. The raw SAM-S phenome (n=1614) significantly underperforms, ranking 6^th^, thus lower than its constituent subgroupings: Shape context (3^rd^, n=300), Zernike moment (4^th^, n=25) and geometrical (5^th^, n=17). It might also be supposed that an imbalance in the number of shape, appearance and motion-associated features in our SAM phenome may bias analysis. However, we do not find evidence of this empirically. The contribution scores computed by SPOT in all studied datasets reflect the expected relative importance of shape, appearance and motion. Notably, SAM-SPOT analysis identified specific shape, appearance and motion-associated modules in a simulated organoid dataset relating to the predefined shape-, appearance- and motion-specific phenotypes (*Reviewers are requested to contact the Editor for access)*.

In contrast, SPOT represents a conceptual workflow specifying a set of standardized processing steps, for which this paper provides an implementation for all steps that is broadly-applicable. However, as in scRNAseq, the actual algorithmic implementation can be customized by users to optimally analyze a dataset-of-interest, when generality is not required. Then, algorithms for each SPOT step should be adjusted to best handle the nuances of the specific dataset, e.g. training custom segmentation and tracking models, training AI-based features instead of the SAM phenome, or alternative dimensionality reduction methods such as t-SNE^99^ or PaCMAP^100^.

Deep learning (DL) AI models with the capacity to learn directly from large input datasets have revolutionized computer vision, delivering state-of-the-art performance in many applications^57,59,101–104^. Consequently, there is much research interest in their application for biological discovery^56^. Here, we compared SPOT analysis with SAM phenome vs AI-based features generated by autoencoder-based models trained by self-supervised learning. We cannot feasibly compare all literature-proposed architectures. Thus, we used a U-Net^105,106^ model, one each for fixed and live-imaging, that is representative of the essential architectural components of the most prevalent family of models used in biological studies: CNN-based autoencoders^47,49,89,107^. Moreover, we trained the models using a training objective of data reconstruction, which makes minimal assumptions of the dataset. Whilst SAM phenomes delivered rich insight for both LIVECell and nuclei division datasets (Fig. 6,7), the neural network models learnt only superficial information and could not identify subtle phenotypes. The SAM phenome also trained superior machine learning classifiers of cell line and mitosis stage than the neural networks, despite for the latter, allowing integrating information from a time window of 5 frames, double the 2 frames used for computing our SAM phenome’s motion features. This performance gap is not simply due to a lack of training data. Indeed, the classification gap was widest in the >1.6 million cell LIVECell. Our results highlight that we cannot assume any AI model to automatically generate the ‘optimal’ features for an analysis. Due to these models being overparameterized and have complex, hierarchical architectures, they can ‘shortcut’ learning only superficial features without additional constraints^108^, and are notoriously difficult to interpret which can lead them to produce unpredictable results^109–111^. Thus, in place of traditional feature engineering, most research efforts in developing performant AI models involves optimizing a complex set of considerations including initial careful dataset curation to ensure diverse and distinct training examples^57,112,113^, and explicitly designing model architectures and training objectives to be most-relevant for the specific downstream analysis^104,112^. Most evident is the unprecedented success of large language models, founded upon switching from CNN-based^105,106^ to more performant but also more resource-intensive transformer-based architectures^114,115^, and more crucially, developing more challenging training objectives such as contrastive learning^116,117^. In biology, recent studies demonstrate the necessity of customized models to obtain features with desired properties such as rotational invariance^49,89^. In short, an AI model capable of generating features comparable to the SAM phenome can only be guaranteed by explicitly designing a custom model with SAM-aware architecture. Lastly, feature interpretation is critical for biological discovery. AI model-features are not suitable for direct interpretation^111^. Further analysis is necessary such as correlating individual features to explicit SAM measurements like area^47,118^. Most detrimentally, being data-driven, AI models trained separately on different unique datasets necessarily yield different, unique feature sets, even if we fix the number of features, i.e., feature 1 in dataset 1 is not the same feature 1 in dataset 2. In contrast, features in our SAM phenome are dataset-agnostic, avoiding any need to retain large raw libraries for retraining. Feature 1 will always be ‘maximum curvature’. Moreover, by design, each feature can be associated with a distinct concept (shape, appearance or motion) and scope (global, regional, pixel-based).

In summary, we have shown novelly that a single set of rationally-designed imaging features can achieve remarkable off-the-shelf performance universally across diverse datasets, while maintaining interpretability not yet easily achievable without significant investment in designing and training custom deep learning models. Together with SPOT, our complete workflow effectively democratizes the complexities of video analysis to the wider scientific and technology community, enabling the detection and characterization of dynamic imaging phenotypes as readily as genotypes in scRNA-seq.

SPOT is currently implemented for 2D videos but this is not a critical limitation. In a companion paper, we demonstrate that SPOT can accurately and comprehensively characterize 3D organoids from 2D projections (*Reviewers are requested to contact the Editor for access*). SPOT can be universally applied to analyze both fluorescently-labelled and label-free live-cell imaging. We provide SPOT as a Python package, which includes computation of the SAM phenome with only standard library dependencies to facilitate customization and extension. For example, users may further augment the SAM phenome to include additional features that are SAM-based or from different modalities e.g. genetics-based, computed by other external software.

## Acknowledgements

This work is mainly funded by the Ludwig Institute for Cancer Research (LICR). FYZ, ANS, XL were funded by the LICR. FYZ was also funded by a EPSRC Life Sciences Interface Doctoral Training Centre EP/F500394/1. HAH, HMB and LM were partly funded by LICR and the Centre for Topological Data Analysis, EPSRC EP/R018472/1. HAH gratefully acknowledges funding from EPSRC EP/R005125/1 and EP/T001968/1, a Royal Society University Research Fellowship (RGF\EA\201074) and UF150238, and Emerson Collective. LM is funded by EPSRC grant EP/R513295/1.

We thank Simon Leedham, Francesco Boccellato, Eric O’Neil, Mary Muers, Kate Dunning, Edward Jenkins, Richard White and Colin Goding for discussion and critical reading of the manuscript.

## Author contributions

Conception: FYZ, XL; Euler characteristic curve shape descriptor development (LM, HB, HH); SPOT and SAM development and analysis (FYZ); GUI interface development and Cellprofiler analysis (ANS). Supervision (XL); Funding acquisition (XL); Writing – Original Draft: (XL, FYZ); Writing – Review and Editing: all authors.

## Resource availability

Further information and requests for resources and reagents should be directed to and will be fulfilled by the lead contact, Xin Lu.

## METHODS

### Construction of 2185 SAM phenome

#### For Shape

we measured 4 area and perimeter statistics to capture size (area, convex area, perimeter and the equivalent circle diameter), 7 shape factor based statistics to capture shape (area-to-perimeter aspect ratio, major axis length, minor axis length, major-to-minor axis length ratio, moment eccentricity, solidity, extent), 7 curvature based statistics to capture the contour complexity after multiplying by the object’s equivalent diameter to make it scale-invariant (maximum, minimum, mean, standard deviation, skew and kurtosis of curvature, and mean absolute curvature), and 5 centroid distance statistics to capture variation of boundary points with respect to cell centroid, as a point of reference (maximum, mean, coefficient of variation, the ratio of maximum-to-minimum distance of a boundary point to the object centroid, and maximum distance between two boundary points) as global features; Hu moments^64^ – a translation, scale and rotation-invariant shape descriptor (total 7 features) based on lower order image moments, Zernike moments^62,65^ – a rotation-invariant shape descriptor based on decomposing the binary shape into its lower-order radial spatial frequencies (total 25 features), and Fourier features^62,63^ – a rotation-invariant shape descriptor based on applying Fourier analysis to decompose the contour into low and high fluctuations, as local-regional features (total 99 features); a histogram of chordal distances (total 8 features with each histogram being a feature), Euler characteristic curve^66^ (ECC) – a topological descriptor based on analyzing the topology of an object at thresholds across a number of directional axes (total 1152 features, considering 36 directions, 32 thresholds), and shape context^68^ – a translation, scale and rotation-invariant descriptor based on capturing the pattern of the contour locally around select ‘keypoint’ boundary points (total 300 features, considering 12 angle bins, 5 distance bins, for 5 equidistantly selected boundary points) as local-distribution features.

#### For Appearance

we measure the mean and standard deviation of intensity values (total 2 features) within the object as global features; the mean and standard deviation of intensity values in each of the three concentric regions, representing an object’s border, middle and inner core, and generated by partitioning the object’s area into 3 bands of equal distance from its boundary (total 6 features) as local-regional features; Haralick features^69^ – a descriptor of local image texture based on measuring statistics of the co-occurrence of a pixel’s intensity value to that of all neighboring pixels within a distance cut-off (total 13 features), and Scale-invariant Feature Transform (SIFT)^71,119,120^ – a translation, scale, and rotation-invariant descriptor of local texture around a keypoint based on equipartitioning the image into 4×4 image patches, and constructing a histogram of the direction of image gradients using 8 angle bins, and concatenating the histograms from all image patches (total 128 features), as local-distribution features.

#### For Motion

we compute first the dense optical flow^121^ for each video using successive frames, giving a velocity vector at each image pixel at a time *t*. We additionally compute the curl and divergence of the optical flow, illustrated in Fig. 1. Curl quantifies the extent of rotational motion around a pixel (positive and negative value for anti-clockwise and clockwise rotation respectively). Divergence quantifies the extent a pixel is a local motion source (positive divergence, local velocity vectors radiate circularly outwards from this pixel) or a motion sink (negative divergence, local velocity vectors radiate circularly inwards into this pixel). We then measure as motion features the mean and standard deviation of the displacement in the object contour between time *t* and *t* + 1 as mean boundary speed statistics (total 2 features), and the mean and standard deviation of the magnitude, curl, and divergence of the optical flow within the object (total 6 features) as global features; the mean, standard deviation, and a 24-bin histogram of the optical flow magnitude in each of the same three concentric bands used for appearance, as local-regional features; SIFT applied to the optical flow magnitude (Speed SIFT, 128 features), curl (Curl SIFT), and divergence (Divergence SIFT) as local-distribution features.

### Validation Datasets

#### Computer vision datasets for testing SAM phenome

Computer vision datasets with manually curated reference annotations were used to validate that our proposed SAM phenome was able to capture and discriminate heterogeneity and be universally applicable as-is without having to refine or define new features. Performance was assessed quantitatively by comparing the result of k-means clustering on the computed respective SAM feature set, where the number of clusters, k was set to the known number of classes in the given dataset. Two standard quality of clustering metrics were used. Adjusted mutual information (AMI, 0-1, *scikit-learn metrics.adjusted_mutual_info_score*) is an adjustment of the Mutual Information (MI) metric which measures the similarity between two clusterings to account for chance, a value of 1 represents perfectly concordant clustering. Adjusted rand index (ARI, −0.5-1, *scikit-learn metrics.adjusted_rand_score*) is an adjustment of the Rand Index metric, which computes a similarity measure between two clusterings by considering all pairs of samples and counting pairs that are assigned in the same or different clusters in the predicted and true clusterings, to account for chance. It is 1 for perfect concordant clustering and lower bounded by −0.5 for particularly discordant clusterings. For the A2D dataset, we also assessed the performance of training a supervised classifier on our SAM features. Performance was reported using the balanced accuracy score (labelled only as ‘accuracy’ in figures), which is the average of the recall obtained on each class to account for imbalanced classes, and F-score (or F1-score), defined as 2*precision*recall/(precision + recall) for a single class. We report the mean across all classes weighted by the number of true instances in each class to account for class imbalance.

#### MPEG-7 Shape Dataset to test shape

The MPEG-7 Core Experiment CE-Shape-1 (shortened to MPEG-7) database^73^ is a standard shape dataset comprising a total of 1400 shapes as binary images; 20 unique image examples each of 70 different shape classes representing different unique everyday objects such as bone, bat, heart, horseshoe, octopus, and turtle, captured with different sizes, rotations, and poses (Fig. 3a). For each image, we binary dilated by 3 pixels, topologically simplified by binary-infilling holes and extracting the external boundary of the shape as a contour of (x,y coordinates) using marching squares (scikit-image *find_contours*). For shapes where the binary-infilling produces multiple disconnected masks (due to a fragmented contour), only the largest connected mask was analyzed. The number of contour coordinates varies across shapes. To ensure the same dimension of SAM phenome is computed, the boundary contour of each shape is resampled to 200 equidistantly spaced boundary points. All SAM-S features were computed and were used to conduct the analysis in Fig. 3 with no other filtering of features except for kernel map dimensionality reduction of ECC features.

#### Brodatz texture database to test appearance

The Brodatz texture database^122^ is a standard dataset for testing appearance descriptors in computer vision. It comprises 112 unique 512×512 pixels images representing different grayscale patterns. Here, we used the normalized Brodatz texture variant^76^, an improved and more challenging dataset that uses an intensity normalization process to eliminate the original’s grayscale background effect. To use the Brodatz textures for testing, we derived a 14,336 image dataset by cropping each of the 512×512 texture images into 64 non-overlapping 64×64 patches and using both the patch and the patch rotated by 90°. SAM-A features were computed directly using the patches, treating the whole patch as an object with a square boundary contour and used to conduct the analysis in Fig. 3 with no other filtering of features. Features were, however, power normalized and z-scored as described for SPOT, Stage 4, step i below.

#### Brodatz textured MPEG-7 shapes to test shape and appearance

To test the combined shape and appearance features, a dataset of normalized Brodatz texture MPEG-7 shapes was created. Five shape classes from MPEG-7 were chosen by equally sampling the measured eccentricity, which was used as a proxy measure of shape complexity. Five normalized 512×512 Brodatz texture images were chosen randomly. To texture the MPEG-7 shapes, their images were all standardized by rescaling to 128×128 pixels. Brodatz texture images were rescaled to 192×192 images. Then for each of the 20 unique images in each MPEG-7 class, we take each 192×192 Brodatz texture image, sample 20 random 128×128 cropped patches and multiply with the MPEG-7 image to create 20 different textured variants of the same basic texture and shape. This gives a total dataset of 10,000 image patches (5 shape classes x 20 images per shape class x 5 texture classes x 20 per texture class). SAM-SA features were obtained by concatenating computed SAM-S (SAM-S (kernel ECC)) and SAM-A features and used to conduct the analysis in Fig. 3 with no other filtering of features. Features were, however, power normalized and z-scored as described for SPOT, Stage 4, step i below.

#### A2D object-motion video dataset to test shape, appearance and motion features in real life conditions

To simultaneously detect joint variations in shape, appearance and motion (SAM) in a setting reminiscent of real application to a heterogeneous dataset we used the A2D dataset^78^. This dataset comprises 3,782 videos sourced from YouTube, depicting seven classes of moving objects (adult, baby, bird, cat, dog, ball and car) performing nine different movements (still, labelled for no movement), climbing, crawling, eating, flying, jumping, rolling, running, and walking (Fig. 4a). No object performs all actions. There are 43 unique object-movement pairs, the frequency of which are unequally distributed. A video may contain more than one instance of an object-movement pair or instances of different object-movement pairs^78^. For each video, 3-5 non-contiguous frames are annotated.

#### Generating consecutive frame object segmentations for A2D dataset to compute SAM phenomes

To compute the full SAM features for each annotated object instance, we first use the Segment Anything model^57^ and the bounding box of its annotated contour to re-segment the object. Then, we used optical flow^121^ to predict the object’s bounding box in the immediate next frame and use it as the prompt to segment its outline using the Segment Anything model. This ensures objects were segmented in the same way across frames.

#### Evaluating clustering and classification performance on A2D test dataset split

The A2D dataset has 43 unique object-movement pairs and is pre-split into 3036 training videos and 746 testing videos. For both clustering and classification analysis, we use object instances in the training videos to set the parameters of the k-means and classifiers, reporting performance only on objects in the testing videos. Full SAM features are defined by concatenating computed SAM-S (SAM-S (kernel ECC), n=562), SAM-A (n=149) and SAM-M (n=422) features and was used to conduct the analysis in Fig. 4 with no other filtering of features. Features were, however, power normalized and z-scored as described for SPOT, Stage 4, step i below.

#### A2D object and movement type similarity graph analysis from UMAP median coordinate

Using the median 2D-UMAP coordinates of each object, movement or object-movement type, compute the pairwise affinity matrix between types defined as where *D* is the pairwise Eucliean distance matrix between coordinates and µ(*D*) is the mean value of the *D* matrix. The final similarity graph plots a connection between type *i* and type *j* if the affinity between types *i* and *j* is greater than the mean value of the affinity matrix (after excluding the main diagonal self-connections).

### Biological cell datasets for testing SPOT analysis

#### LIVECell dataset of diverse cell morphologies to test SPOT analysis

The LIVECell^86^ dataset consists of 5,239 manually annotated, expert-validated, Incucyte HD phase-contrast microscopy images with a total of 1,686,352 individual cells annotated from eight different cell lines: A172 (glioblastoma), BT-474 (ductal breast carcinoma), BV2 (microglial mouse cells), Huh7 (hepatocyte), MCF7 (human breast cancer), SH-SY5Y (neuroblastoma), SKBr3 (human breast cancer), and SK-OV-3 (ovarian cancer). The images and corresponding cell masks are split into one train-val folder of 3,727 images and a test folder of 1,512 images. We used only the test folder images for SPOT analysis. The provided cell masks were used as-is without checking for and removing potential partial cells on the image borders prior to SAM feature computation. As cells are fixed, only shape and appearance features were measured. The compiled SA-phenomes were filtered and processed as described below but without performing those preprocessing steps specific for videos where time information is available and SPOT analysis (Stage 4) was applied. For comparing the training of a classifier based on SPOT SAM or neural network features, we used the train-val folder images to train the classifier and the test folder images to evaluate performance.

### Single-cell tracking datasets to test SPOT temporal analysis

We selected two datasets (U373 and HL60) from the 2D single cell tracking challenge^98^ with distinct and easily interpretable phenotypes (distinct cell morphologies as U373 cells migrate, and distinct appearance changes as HL60 cells divide) to test SPOT analysis (Stage 4). Each dataset was comprised of two unique videos, with each video’s outline annotation and temporal tracking information already provided for all timepoints, which we use to avoid introducing additional errors in segmentation or tracking in SPOT, Stage 3. Full SAM features were computed and compiled for all annotated cell instances that were fully present in the imaged field-of-view. Cells on the border which were partially in-view were removed prior to SAM feature computation and the corresponding single cell tracks truncated correspondingly into separate contiguous tracklets. The compiled SAM phenomes were filtered and processed as described below and SPOT analysis (Stage 4) was applied to both datasets in the same manner, using all the data instead of setting a percentile cutoff to exclude potential outliers when computing the representative phenotype cluster exemplar images.

### HeLa cell histone H2B dataset to test SPOT temporal analysis

We use a published dataset of 7 timelapse microscopy movies of human tissue culture cells (HeLa ‘Kyoto’ cells) expressing a fluorescent chromatin marker (histone H2B–monomeric (m) Cherry)^90^. The movies comprise 326 video sequences, whereby each sequence tracks a unique single cell, showing only its nucleus and the change in nucleus morphology as the cell undergoes interphase and the five phases of mitosis (prophase, prometaphase, metaphase, anaphase and telophase). For each frame the mitosis stage is provided but not the nuclei segmentation. To compute SAM features, we custom segmented the nuclei frame-by-frame by binary intensity thresholding. Briefly, each image was contrast enhanced by clipping the intensity to the 2^nd^ and 99^th^ percentile; subject to adaptive histogram equalization (*skimage.exposure.equalize_adapthist*) with a clip limit of 0.02; and median filtered with a size of 3 pixels. The image was binary Otsu thresholded to generate the segmentation which was then postprocessed by morphological binary closing (disk kernel, radius=3 pixels) and binary filling holes. Connected component analysis was then applied to retain the spatial component closest to the image centroid. To correct for, particularly in anaphase, two separated chromosomes being erroneously segmented as one, watershed segmentation was applied for separation, only the most centrally located chromosome was retained in the image for SAM feature computation. Full SAM features were computed for each segmentation. The compiled SAM phenomes were filtered and processed as described below and SPOT temporal analysis (Stage 4) was applied.

### CellProfiler features computation, processing and comparison with SPOT

#### Computation of CellProfiler shape-related metrics

All 1400 image files in the MPEG-7 dataset were uploaded to CellProfiler v4.2.6. The images were processed using CellProfiler’s ‘ConvertImageToObject’ and ‘MaskObject’ modules in succession. The ‘MeasureObjectSizeShape’ module was then used to generate shape-related metrics derived from object boundaries. These metrics were exported using the ‘ExportToSpreadsheet’ module, before being filtered to remove irrelevant columns (such as centroid measurements) and non-numerical columns prior to analysis.

#### Computation of CellProfiler appearance-related metrics

All 22,400 images in the data augmented Normalized Brodatz dataset were uploaded to CellProfiler v4.2.6 (112 Brodatz image classes x 200 patches per Brodatz class). Generation of appearance-related metrics was completed by grouping the images into 10 batches to prevent crashing. Within each batch, patches were grouped based on their image of origin. The ‘MeasureGranularity’, ‘MeasureTexture’, ‘MeasureImageIntensity’, ‘MeasureImageQuality’, and ‘MeasureImageSkeleton’ modules were run in succession, selecting for all metrics available to be computed within each module’s settings, wherever relevant. These metrics were exported using the ‘ExportToSpreadsheet’ module, and filtered to remove duplicated, irrelevant (such as execution time-related), and non-numerical columns prior to analysis.

#### Computation of CellProfiler shape and appearance-related metrics

The 10,000 images in the Brodatz textured MPEG-7 shapes dataset we created were cropped using the tightest bounding box around the shape, then uploaded to CellProfiler v2.4.6. The appearance-related metrics were computed at the whole-image level for each crop, a design choice made due to inadequate performance of the ‘ConvertImageToObject’ module on this dataset. The shape-related metrics are identical to those calculated for the entire MPEG-7 dataset and thus were not recomputed. All appearance-related metrics were computed by running the ‘MeasureGranularity’, ‘MeasureTexture’, ‘MeasureImageIntensity’, ‘MeasureImageQuality’, and ‘MeasureImageSkeleton’ modules in succession. These metrics were exported using the ‘ExportToSpreadsheet’ module, and filtered to remove duplicate, irrelevant, and non-numerical columns prior to analysis. The shape-related metrics and appearance-related metrics were concatenated to form the combined CellProfiler shape-appearance descriptor.

#### Analysis of CellProfiler computed features

To ensure like-for-like comparison, CellProfiler-computed features were handled and processed in the exact same manner as SPOT’s SAM features. This was the case for each relevant validation dataset.

### CNN autoencoder features computation, processing and SPOT analysis

#### Training Data

Individual cells of the LIVECell dataset were cropped using the bounding box of their segmentation mask, defined by upper left corner coordinates, (*x*_1_,*y*_1_) and lower right corner coordinates (*x*_2_,*y*_2_) where *x*_1_,*x*_2_ is the minimum and maximum value of the x-coordinate, and *y*_1_,*y*_2_ is the minimum and maximum value of the y-coordinate across all pixels comprising its segmentation. Cropped images were resized to a standard size of 64 x 64 pixels, the same size used by SPOT to compute SAM features. Image intensities were min-max normalized per image to be in the range 0-1. Cells from the train-val dataset were used for CNN autoencoder training, reserving the test dataset for SPOT analysis and comparison with SAM-SPOT.

#### Model Training

A convolutional neural network (CNN) was trained using Keras with Tensorflow backend with the Adam optimizer (default parameters) for 100 epochs, a batch size of 128 images and mean absolute error loss. The encoder network uses 4 convolutional 2D layers with 3×3 kernel, downsampling by stride 2 between layers. The first layer uses 32 filters, with the filter number doubling for each subsequent layer. To compute the 256 CNN features, the encoder output is flattened, processed by a dense layer of 1024 units, then by a dense layer of 256 units. The decoder network mirrors the encoder in reverse, transforming the 256 CNN features back to an image output. ReLU activation was used for all layers throughout.

#### SPOT Analysis of CNN autoencoder features

The 256 CNN features per image were used as-is with no other preprocessing. The same SPOT analysis with identical parameters as for SAM-SPOT was performed using the CNN features.

### CNN-LSTM autoencoder features computation, processing and SPOT analysis

#### Training Data

The HeLa cell histone H2B dataset^90^ provides already cropped and centered nuclei videos for analysis. We resize individual video frames to a standard size of 64 x 64 pixels, the same size used by SPOT to compute SAM features. Image intensities were min-max normalized per image to be in the range 0-1. The individual cell videos were processed to generate all possible continuous 5-frame image sequences, that is frames 1 to 5 is one sequence, 2 to 6 another, 3 to 7 another etc (Suppl. Fig. 8b).

#### Model Training

The convolutional neural network (CNN) architecture used for the CNN autoencoder analysis of the LIVECell dataset was adapted to incorporate temporal video information into model training. We apply the same CNN encoder to each frame of the input 5-frame sequence, process the outputs using a bidirectional LSTM layer of 16 filters to incorporate time dependencies, and then apply a dense layer of 256 units followed by the same CNN decoder to each frame to transform outputs back to images. This CNN-LSTM model was trained using Keras with Tensorflow backend for 100 epochs with a batch size 128 sequences and mean absolute error loss. After training, the CNN encoder can be used to extract the 256 CNN-LSTM features for single timepoint images.

#### SPOT Analysis of CNN-LSTM autoencoder features

The 256 CNN-LSTM features per image were used as-is with no other preprocessing. The same SPOT analysis with identical parameters as for SAM-SPOT was then performed using the CNN-LSTM features.

### Assessment of classification performance using SPOT SAM and CNN autoencoder features

#### LIVECell dataset^86^

Due to the large size of this dataset, we trained a two-layer multilayer perceptron (MLP) classifier with categorical crossentropy loss to predict the cell line from input features in Keras with Tensorflow backend. We weight the loss inverse to the number of cells in each cell line to account for the discrepancy. Each layer used 100 units and tanh activation. The Adam optimizer (learning rate = 5 × 10^-4^, p_1_ = 0.5) was used to train the minimum validation loss model with early stopping (patience=5, min_delta = 0.0005). We train the optimal classifer for the test folder for both SPOT SAM and CNN autoencoder features by using the test folder as validation set for early stopping. We use the preprocessed SPOT SAM features (709 features) for training the classifier. All dataset dependent preprocessing steps such as zero-valued and high variance features, kernel transform dimensionality reduction of ECC shape features, standard scaling and power transform normalization was set based on the train-val folder images and then applied to the test folder images to ensure features were processed in the same manner. As the CNN autoencoder features (256 features) were learned from images directly, we used them as-is to train the classifier without any preprocessing.

#### HeLa cell histone H2B dataset^90^

The total number of nuclei is relatively small compared to LIVECell (12,714 = 326 video sequences x (40-1) frames), therefore we trained a support vector machine (SVM) classifier (using the scikit-learn library: RBF kernel, C=0.1, random_state=0, class_weight=’balanced’, decision_function_shape=’ovr’) to predict interphase and the five mitosis stages (prophase, prometaphase, metaphase, anaphase and telophase) from inputfeatures. An MLP-trained classifier was also trained but was worse than RBF SVM (data not shown). The video sequences were evenly split according to the 7 videos they originate from to construct training (videos ‘0037’, ‘0039’, ‘0042’, ‘0046’) and testing (videos ‘0038’, ‘0040’, ‘0045’) video frames. Classifier performance was reported only using the testing video frames. We use the preprocessed SAM features (186 features) used in SPOT analysis to train the SVM classifier. We use the CNN-LSTM autoencoder features (256 features) learnt from video frames directly, as-is to train the SVM classifier without any preprocessing.

### Determining the optimal number of boundary points used to represent 2D shapes

We used the diverse shapes of the MPEG-7 dataset to determine the optimal number of boundary points. For each object, we use the contour extracted using marching cubes (scikit-image *find_contours*) as described above as its ground-truth reference boundary. Then, for each object, we resampled its reference boundary with *n* equidistantly spaced boundary points, increasing *n* from 10 to 1000 at increments of 50. The directed Hausdorff distance, *H(A,B)* measures the largest of all distances from a point in a point set, *A* to the closest point in another point set, *B*. In general, *H(A,B) * H(B,A)*. We therefore defined the shape reconstruction error of the *n* boundary point resampled contour as the maximum of its two directed Hausdorff distances (*max(H(A,B),H(B,A)))* with the reference boundary. Our choice of *n* = 200 equidistance boundary points balances thisreconstruction error with using a minimal number of points to minimize hard disk storage (Suppl. Fig. 1).

### Shape, appearance and motion Phenotype Observation Tool (SPOT)

SPOT sets out several stages for analysis as depicted in Fig. 2.

#### Stage 1: Preprocess video datasets

SPOT requires 2D videos without stage drift or other image translation artefacts. To ensure this, raw acquired microscopy videos are preprocessed. Preprocessing involves (i) if applicable, the projection of any 3D video frames where each frame is acquired as a 3D z-stack to single 2D images. As default, we use extended-focus projection for label-free microscopy imaging and maximum intensity projection for fluorescent imaging, to optimally retain object features and maximize the number of objects visualized in the 2D projection image. 2D videos are then (ii) frame-by-frame registered to remove any potential global translation motion. We do this by firstly detecting and removing any blurry frames, automatically detected as having a mean image intensity gradient magnitude one standard deviation less than the mean computed across all video frames. Fourier transform cross-correlation^123^ was then applied sequentially to consecutive frames to register the entire video. Registration is essential for extended long-time timelapse data acquired in multiple acquisitions which often shifts the imaging position, sometimes by a significant distance.

#### Stage 2: Detect video objects

Individual object instances must be segmented from video frames to compute Shape and Appearance features. To also compute all motion features, every object must be tracked to link an object instance with the matching instance in the next frame. Tracking is also essential to identify and remove any transient objects whose tracks only last a few frames. In this study, we used, unless otherwise described, the provided segmentations and object tracking of each public dataset, if available.

For general use, the SPOT code repository only provides a custom trained organoid bounding box detector and segmentation model, developed for our companion organoid study but no general segmentation models, as segmentation is often more optimal, if fine-tuned to individual datasets and the specifics of video acquisition. We note general segmentation models for 2D images of both natural objects^57^ and cells^124^ exist, and readily obtained from their respective sources. SPOT does however provide a general tracker to track segmentations provided as bounding boxes based on spatial overlap across successive frames to generate temporally consistent tracklets. Notably, linking between consecutive frames is aided by using the computed dense optical flow to handle potential large shape, appearance and movement changes. Tracklets are built starting with each detected box in frame 0. The coordinates of each bounding box in the next frame, (*x*(*t*+ 1),*y*(*t* + 1)), is predicted by fitting a rigid motion model (scale, translation, rotation), *i* to the local optical flow, (Δ*x*,Δ*y*) within the box, i.e. *(x(t+ 1),y(t+ 1) = (x(t) + Δx(t),y(t) + Δy(t) = i((x(t),y(t))*. Using the predicted coordinates after fitting i((x(t),y(t))), the boxes are matched to actual boxes of detected objects in the next frame, frame 1, via linear assignment bipartite matching (c.f. Scipy *linear_sum_assignment*), using the pairwise cost matrix constructed from 1-intersection-over-union (IoU) between predicted and actual detected boxes in frame 1.

A minimal IoU cut-off is used to determine a successful pairing (default IoU≥0.25, i.e. 1-IoU< 0.75). The tracklets of successfully matched boxes are extended. Tracklets with unsuccessfully matched boxes in frame 0 are also extended opportunistically using the motion model predicted coordinates. Boxes unsuccessfully matched in frame 1 start new tracklets, adding to the pool of running tracklets to extend in subsequent frames. The linking process is repeated frame-by-frame consecutively until the end of the video and all detected boxes are a member of a tracklet. Running tracklets that have not been successfully matched to a ‘real’ object box corresponding to a segmentation after a specified number of frames are ‘terminated’ and no longer considered. This tracking process is designed to ensure coverage of all segmentations throughout the length of the video. Tracklets can finally be filtered by setting threshold values for 1) minimal lifetime coverage (measured as a fraction of tracklet length) and 2) mean confidence detection score across all boxes composing the tracklet (e.g. confidence score predicted by object detector or segmentation probability map).

#### Stage 3: Computation of shape, appearance, and motion (SAM) features

SAM features were computed with custom code using, where possible, standard implementations available in established Python libraries such as scikit-image and OpenCV (see our GitHub code, https://github.com/fyz11/SPOT). Detailed descriptions and relevant references for each feature are provided in Suppl. Table 1. Motion features are only computable for an object instance at timepoint *t* if it has a corresponding instance in thenext timepoint, *t* + 1. Most features in our SAM phenome are dimensionless (Suppl. Table 1). Where a measurement is not dimensionless, all measurements were extracted in standard physical units of micrometer (μm) for length and hours (h) for time.

#### Stage 4: Temporal analysis of shape, appearance, and motion (SAM) features

This stage comprises two parts which can be performed independently and then mix-and-matched. Part A implements a streamlined workflow to monitor temporal changes in phenotypic heterogeneity. Two primary analyses are performed: (1) a SAM population phenotype trajectory providing a continuous readout of temporal evolution driven by majority phenotypic states and (2) automatically finding a finite number of discrete phenotype clusters to characterize the full phenotypic heterogeneity, and the temporal phenotype cluster composition and inter-cluster transitions within a condition provide further interpretation of its phenotype trajectory. Part B focuses on analysis interpretation by capturing the Pearson correlation between individual SAM features across all objects in the analyzed dataset to consolidate individual features into a smaller number of ‘SAM modules’, similar to molecular pathways. The SAM modules is used to retrieve exemplar object images from the dataset, and find the driving features for interpretation.

### Part A: Discovery and analysis of dynamic phenotypic heterogeneity Step i*: Generating filtered SAM phenomes for analysis*

Raw computed SAM phenomes require filtering; to remove those features that are too small or zero-valued and those that exhibit too high a variation across the dataset to be analyzed. Specifically, we remove all features that are zero valued and those with zero variance across all objects (from possibly different experiments) to be compared in the same analysis. High variance features (defined typically >2 standard deviations of the global mean) are also removed. The remaining features are normalized so that the value of each feature is comparable across features (for example, to enable shape area to be considered with equal weighting to shape curvature, where the raw values differ greatly in magnitude). First, all curvature-associated features with units of length^-1^ are multiplied by the equivalent diameter to correct for the inherent bias of measuring decreased curvature for objects of the same shape but of different size. Second, a radial basis function (RBF) kernel is applied using the Nystroem approximation, (c.f. scikit-learn *Nystroem*) to compress the raw, sparse and discrete Euler characteristic curve (ECC) features to 100 dimensions which we found yielded improved clustering performance and made the inclusion of ECC features more beneficial than its exclusion (see Suppl. Table 2, SAM-S (ECC) vs SAM-S (kernel ECC)). Third, for data acquired as separate batches, we correct feature values to remove the ‘batch’ variation using linear regression. We consider ‘batch’ as the primary experimental source of variation, for example, a ‘batch’ could be individual plates imaged on different days or different patients. Fourth, the (possibly batch corrected) feature values are normalized by applying standard scaling (scikit-learn *StandardScaler*) followed by a power transformation (scikit-learn *PowerTransformer*, method=’yeo-johnson). The normalized feature values or SAM scores are plotted as ‘Score’ in figures. For timelapse imaging, particularly for label-free imaging, ridge regression (scikit-learn *Ridge*, alpha=1) using the frame number as the dependent variable and the SAM score of each feature as independent variables is used to select only those features that exhibit significant temporal variation - those with an absolute regression coefficient value greater than the mean absolute regression coefficient value across all features.

### Step ii: Low-dimensional embedding to construct the SAM phenomic landscape

The SPOT SAM phenomic landscape aims to capture and visualize all SAM-based phenotypic variations across every object, every timepoint and every video in a single 2D coordinate space. As such, though any dimensionality reduction can be applied (Suppl. Fig. 3), for best results the dimensionality reduction method should ‘spread’ the SAM variation in 2D space instead of collapsing object instances into branches or singular points as in scRNAseq for lineage inference. Consequently, we chose to use Uniform Maniform Approximation and Projection (UMAP)^125^ developed based on mathematical theory of topological spaces via the Python *umap-learn* implementation to project preprocessed SAM features into 2 dimensions for plotting and analysis. The following default parameters are used: n_neighbors=100, random_state=0, spread=0.5, min_dist=0.5, metric=’euclidean’. For very large datasets (>1 million points), or if it is required to be able to add new object points after initial mapping, we suggest users instead use a dimensionality reduction method supporting batch-based evaluation such as parametric UMAP or deep neural autoencoders.

### Step iii) Computation of the SPOT SAM population phenotype trajectory

To construct temporal SAM phenotype trajectories in the phenomic landscape, the total temporal duration of video acquisition was partitioned into equal time intervals. For each time interval, a heatmap density image (see below) was computed of only those individual object instances present during that time interval. The heatmap image was partitioned into high- and low-density regions using the mean density (± 2 or 3 standard deviations) as a cut-off. The mean UMAP coordinate of the high-density regions (possibly more than one) is computed as the phenomic landscape coordinate that best summarizes the distribution of majority phenotypes during this time interval. The full phenotype trajectory is formed by chronologically linking together the mean UMAP coordinates of high-density regions from all time intervals.

### Step iv: Partitioning of the phenomic landscape into discrete phenotype clusters

Clustering is used to mutually exclusively partition the phenomic landscape into a discrete number of phenotype clusters such that the object instances within each phenotype cluster share similar SAM features. Here we use k-means clustering to generate approximately equal-sized phenotype clusters. This equates to using a noninformative prior of the occurrence of phenotypic states. If it is known or hypothesized or desired that phenotype clusters should have different abundances, for example, we expect rare phenotypes, then alternative clustering such as density-based clustering methods may be more suitable and can be used instead. Scikit-learn k-means clustering (random_state=0) was applied to the UMAP projected 2-dimensional coordinates to cluster individual object datapoints into a discrete number of ‘phenotype clusters’ representing a set of prototype SAM characteristics for the dataset. As such, these clusters do not necessarily represent only biological phenotypes but also capture salient imaging artefacts such as blur or out-of-focus objects. The number of clusters was determined from a minimum of two clusters to a maximum of 20 clusters guided by the minimum of the elbow method applied to the k-means inertia loss. Using these phenotype clusters, the phenotypic diversity of a collection of object instances can be described by a histogram recording the occurrence frequency of each prototype. This approach naturally reconciles multi-parametric phenotypic heterogeneity and is motivated by the success of bag of words/features approaches in computer vision^126^.

#### Auto-generating exemplar object instances for each phenotype cluster

The first principal component (PC) obtained by applying PCA to all object instances within a phenotype cluster provides a measure of the contribution of each object instance to the variance within the cluster. Object instances within a cluster are ranked in descending order of the first PC value and the top rankings are visualized as exemplars and are aligned with the first PC. As PCA is sensitive to outliers caused by errors in the initial object segmentation or tracking, ranking was performed only for instances for which the values of the first PC were less than a specified percentile (for A2D^78^, 90^th^ percentile, for LIVECell^86^, 100^th^ percentile and for the HeLa cell histone H2B dataset^90^, 75^th^ percentile). For the single cell tracking datasets^98^, where both reference segmentations and tracking were provided, we use all object instances without exclusion (i.e. the 100^th^ percentile as cut-off).

### Step v) Monitoring phenotype cluster dynamics

#### Temporal phenotype cluster frequency stacked barplots

Each object instance is assigned the phenotype cluster into which its 2D-UMAP coordinates fall. As for phenotype trajectory computation, the total temporal duration of video acquisition was partitioned into equal time intervals. For each time interval, a normalized histogram of the number of object instances within the interval is constructed where (the frequency of phenotype cluster *i*) = (# object instances in time interval of cluster id *i*) / (total # object instances in that time interval). The normalized histograms for all time intervals are concatenated chronologically to form a stacked barplot of phenotype cluster frequency over time.

#### Hidden Markov Model (HMM) phenotype cluster transition graph

To compute the HMM phenotype cluster transition graph requires the objects-of-interest to be tracked over time. Each object instance is first assigned the phenotype cluster corresponding to its 2D UMAP coordinate. Then, using the object tracks from SPOT, Stage 2, we construct the phenotype cluster label timeseries for each tracked object (Fig. 5c). We temporally model the phenotype cluster label timeseries as individual observed sequences of a hidden first-order Markov process whereby the probability of a phenotype cluster transitioning to another cluster in the next frame is fixed and independent of its historical state. We fitted the phenotype cluster label timeseries using the categoricalHMM model in the Python hmmlearn package. The number of hidden states was specified to be the same as the number of phenotype clusters and the identity of each hidden state was one-to-one matched to a phenotype cluster after HMM fitting, by comparing the HMM predicted and input phenotype cluster label timeseries. The phenotype cluster transition probabilities is then the transition matrix of the fitted HMM. The cluster transition graph visualizes the transition probabilities, with increased arrow opacity representing a higher probability. The graph was generated using the Python hmmviz package. To compute the transition entropy of each phenotype cluster, the Shannon entropy was computed for each row of the cluster transition probability matrix.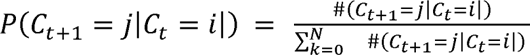

### Part B: Discovery and analysis of dynamic phenotypic heterogeneity

#### Step vi) Automatic grouping of filtered SAM phenome features into SAM modules

SAM modules group features that covary in the dataset, specifically features that exhibit greater Pearson correlation between those features in the same module than between those features in a different module. To identify SAM modules, we use all of the compiled SAM phenomes to be analyzed across all conditions. Following filtering and preprocessing above, (SPOT Stage 4A, step i) the (# features) x (# features) pairwise Pearson correlation coefficient matrix between the retained and transformed SAM features is computed using all object instances as ‘samples’. To automatically group features into SAM modules and determine the number of SAM modules, we use hierarchical clustering with average linkage, together with clustering quality metrics. Hierarchical clustering generates a dendrogram whereby the root node groups all features as one, and progressive branch splitting corresponds to splitting of the larger parental grouping into unique smaller mutually exclusive groupings. At the bottommost leaves of the dendrogram, the individual SAM modules are the individual SAM features. To determine the optimal number of modules, we used as the clustering quality metric, the Davies-Bouldin (DB)^127^ score, defined as the average similarity measure of each cluster with its most similar cluster, where similarity is the ratio of within-cluster to between-cluster distances. The lower the DB score, the better the clustering, with the minimum DB score as 0. Computing the DB score as a function of the increasing number of modules corresponding to progressive dendrogram branch splits, we observe the DB score increases then decreases with typically, multiple peaks. We set the number of modules corresponding to the first peak in the DB score. This strikes a balance between having too few and too many modules to describe phenotypic heterogeneity.

#### Step vii) Finding exemplar object instances and evaluating SAM module expression in individual phenotype clusters

Principal component analysis (PCA) finds a linear transformation of the data such that the first coordinate or first principal component corresponds to the direction of greatest variance of the data. The expression of SAM module *i* for each object instance is defined as the first principal component after applying PCA to the preprocessed and normalized SAM features from SPOT Stage 4A, step i (i.e. the SAM scores of final features) that form SAM module *i*. As the normalized SAM features have zero mean and the same standard deviation, and as PCA without whitening preserves this property for SAM expression, we define a score of the extent an object instance expresses SAM module *i* as the expression of SAM module *i* minus the maximum expression for any other SAM module. For each SAM module, object instances are ranked in descending order of this score and the top instances are visualized as exemplars.

### Scoring the contribution of feature type and spatial scope

Principal component analysis (PCA) finds a linear transformation of the given input data such that the first principal component corresponds to the direction of greatest variance of the data. SPOT uses this property to derive a score of the contribution of feature type; shape, appearance or motion and spatial scope; global, regional or distribution to the phenotypic variation in each dataset. Each feature in the SAM phenome is annotated with its feature type and spatial scope (see Suppl. Table 1). To compute the contribution of feature type, we evaluate first the ‘expression’ of shape, appearance and motion of each object instance, defined by the first principal component after applying PCA to the subset of pre-processed and normalized SAM features from SPOT Stage 4A, step i, categorized as shape, appearance and motion respectively. PCA is applied again to all object instances’ shape, appearance and motion expression but treating the expression as ‘samples’ i.e. rows and object instances as ‘features’ i.e. columns. The absolute value of the coefficients of the first principal axis, associated with the first principal component, defines the contribution score of shape, appearance, and motion, respectively, to the data variance. The contribution of spatial scope is computed separately in the same manner.

### Phenomic landscape density heatmaps

To produce flow cytometry-like density heatmaps of 2D UMAP projected points of object instances efficiently, we estimate the Gaussian kernel density of points using an image-based approximation. We rasterize the UMAP plot as an image of desired resolution (e.g. 2000 x 2000 pixels). Individual 2D UMAP coordinates *(umap_X_, umap_y_)* are transformed to integer image coordinates using 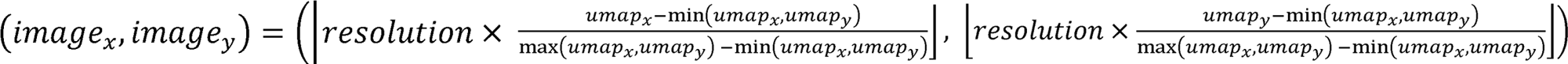 where [] is the floor operator, max is the maximum value of UMAP *(x,y)* coordinates plus a padding value (we use 1) and min is the minimum of UMAP *(x,y)* coordinates minus a padding value (we use 1). Then, each UMAP point for the condition under consideration contributes +1 to the image pixel value at the computed corresponding image coordinate, producing a count of the number of individual points that map to a particular image pixel. The resultant image is finally Gaussian smoothed with an automatically determined sigma, 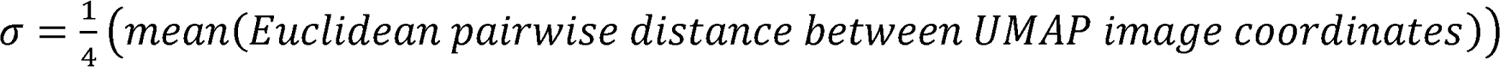. The full pairwise distance matrix between all points does not need to be used for computation. We found a random sample of 1000 points sufficient to obtain a smoothly varying density heatmap.

### Clustering of SAM temporal trajectories

To cluster the temporal phenotype trajectories (computed as described above) of objects from different conditions, a pairwise distance matrix is constructed using multidimensional dynamical time warping (DTW) (through the *dtaidistance* Python package) to measure the distance between any two trajectories. Compared to pairwise Euclidean distances, the use of DTW accounts for variations in trajectory speed. The resulting distance matrix, *D*, is unbounded in magnitude; for clustering *D* is transformed into an affinity matrix, *A* with entries in the range from 0-1 using the formula, 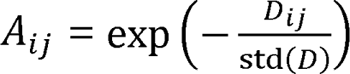 where std() is the standard deviation of distances in *n*. The rows of the affinity matrix are used as the features describing the trajectory of each condition. By default, the average linkage and the Euclidean metric were then used to hierarchically cluster different conditions by their features.

## Data and code availability

The developed general SPOT Python library used to compute and analyze SAM phenomes is available at GitHub, https://github.com/fyz11/SPOT for free academic use with example scripts and example data. Any additional information required to reanalyze the data reported in this paper is available from the authors upon suitable request.

---

**Supplementary Figure 1.**
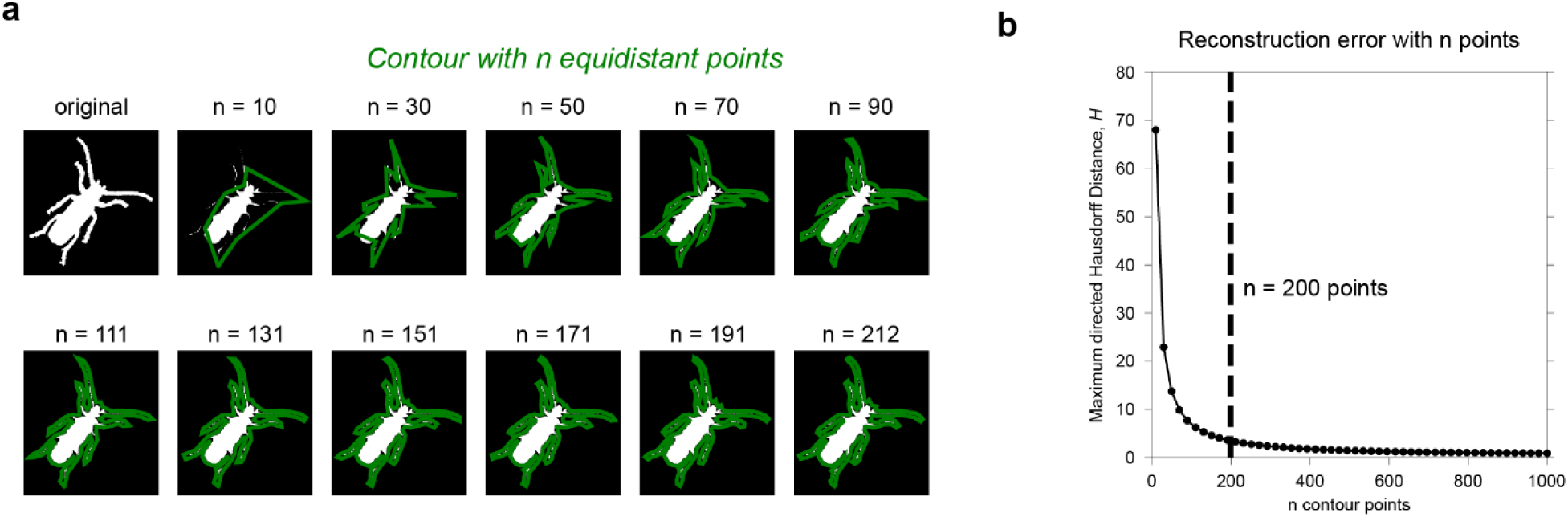
Assessment of the number of equidistantly sampled contour points to represent a shape contour. a) Resulting contour (green) of a beetle image in the MPEG-7 dataset when we use *D* points equidistantly sampled from the true contour. As *n* increases, the more accurate the approximation. **b).** Mean of the maximum directed Hausdorff distance, *H* between the *n*-point contour approximation and the true contour over all images in the MPEG-7 dataset, as *n* is increased. The dashed line shows the *n* = 200 used as default in SPOT which achieves a balance between minimizing the number of points and the reconstruction error.

**Supplementary Figure 2a.**
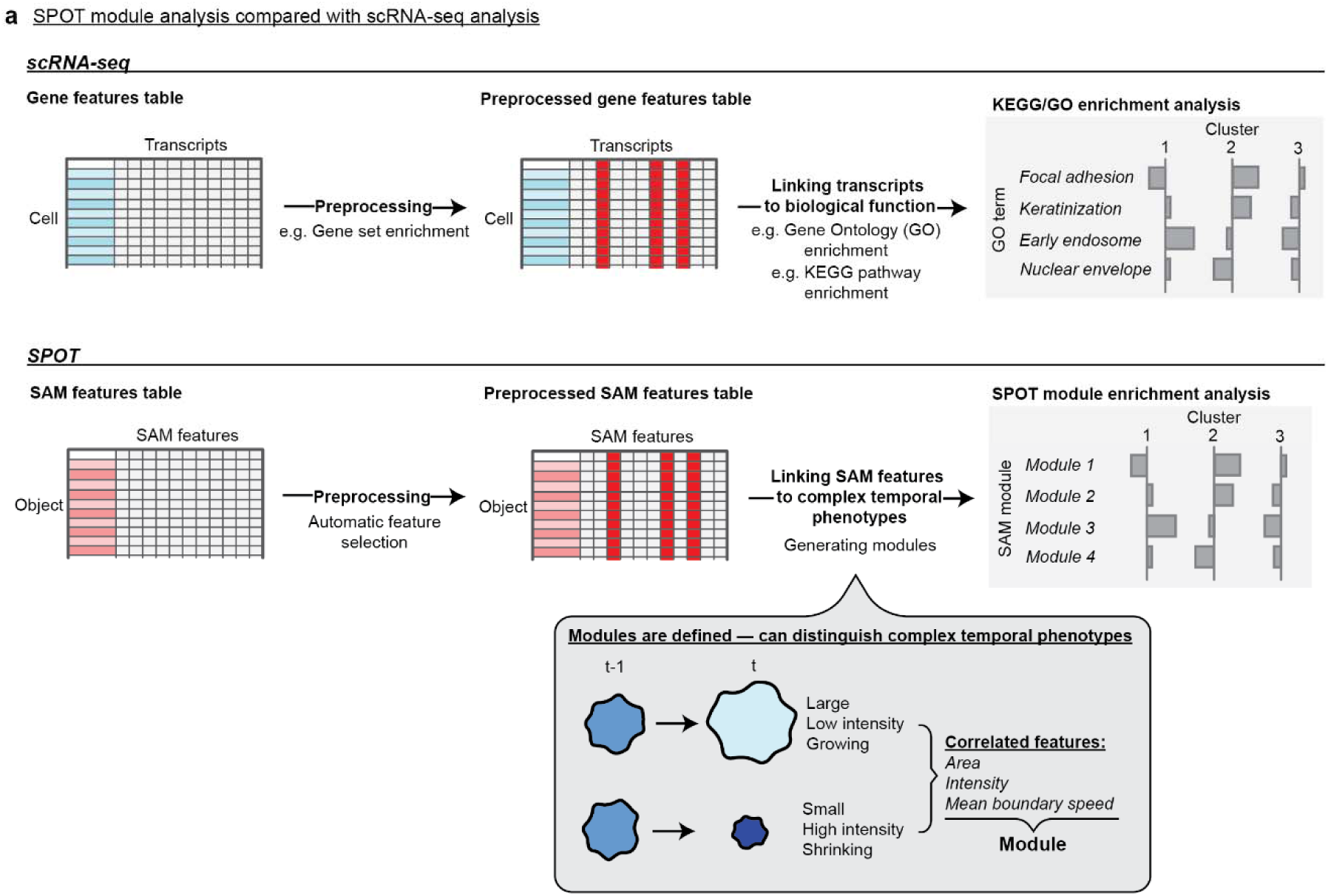
SPOT module analysis is analogous to enrichment analysis after scRNA-seq. In scRNA-seq analysis, the expression scores of the filtered and preprocessed transcripts are used to identify enrichment of biological pathways via gene ontology (GO) or KEGG pathway enrichment. Similarly, SPOT module analysis hierarchically clusters features in the filtered and preprocessed SAM phenome to first identify modules and then perform module enrichment. Each module represents a more conceptually higher-level complex phenotype than any single feature. Module enrichment analysis scores the ‘expression’ level of each module within a given grouping of object instances.

**Supplementary Figure 2b.**
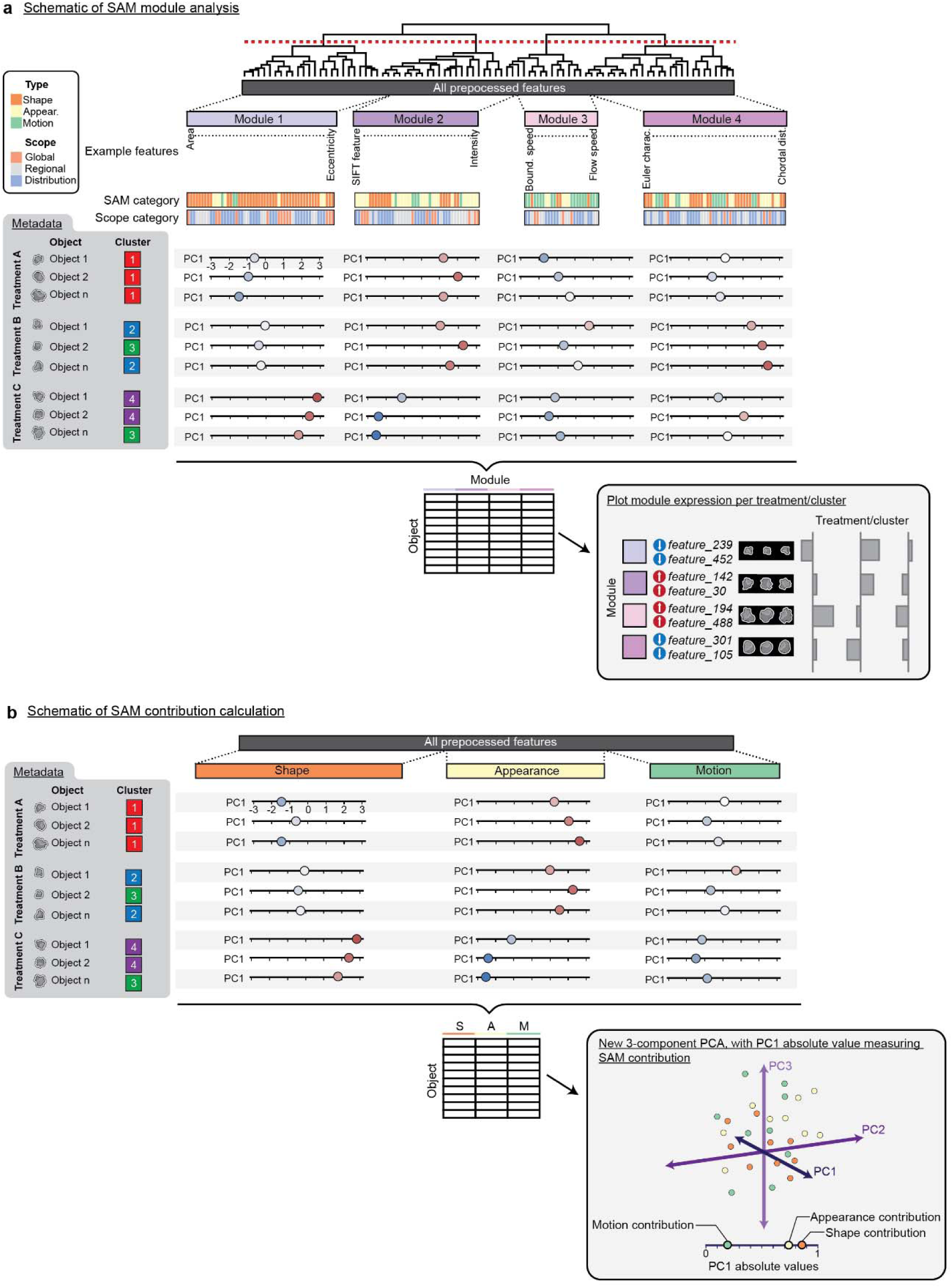
Schematic of SPOT module analysis and computation of SAM/scope contributions. **a)** Identification of SAM modules, groupings of similarly covarying single SAM features by hierarchical clustering. Example SAM and scope categories are displayed for each module, illustrating that correlated features within a module comprise diverse SAM and scope categorizations. A module represents a complex SAM phenotype that cannot be captured by any single feature. For each module, an expression score is generated by calculating the first principal component (PC1) score for each object instance (Methods). The PC1 score measures the contribution of an object instance to the axis of maximal variance, so a high positive PC1 score for a given object instance indicates that its module feature values align closely with PC1. The PC1 expression score can be both positive and negative. A high positive expression indicates the module contributes significantly to the overall variance of the object instances in the dataset, and an enrichment of the complex phenotype the module represents. A high negative expression also indicates a significant contribution of the module, but the object instances in the dataset display the inverse of the complex phenotype the module represents e.g. if the module represents an elongated shape phenotype, the inverse phenotype is a spherical shape. Individual modules are interpreted by finding representative exemplar object instances and top driving single features (Methods). Module expression can be computed for any grouping of object instances by experimental condition, timepoint, or SAM phenotype cluster for comparative analysis. **b)** Shape, appearance, and motion contribution scores can be generated by individually subdividing the features into their SAM categories which serve as modules, so that for each category, we compute the PC1 scores as the category expression for all object instances. A PCA is performed with the category expression to determine the contribution of each category to the first principal component axis. The absolute value of the contribution is defined as the contribution of the SAM category on a scale of 0 to 1, (Methods).

**Supplementary Figure 3.**
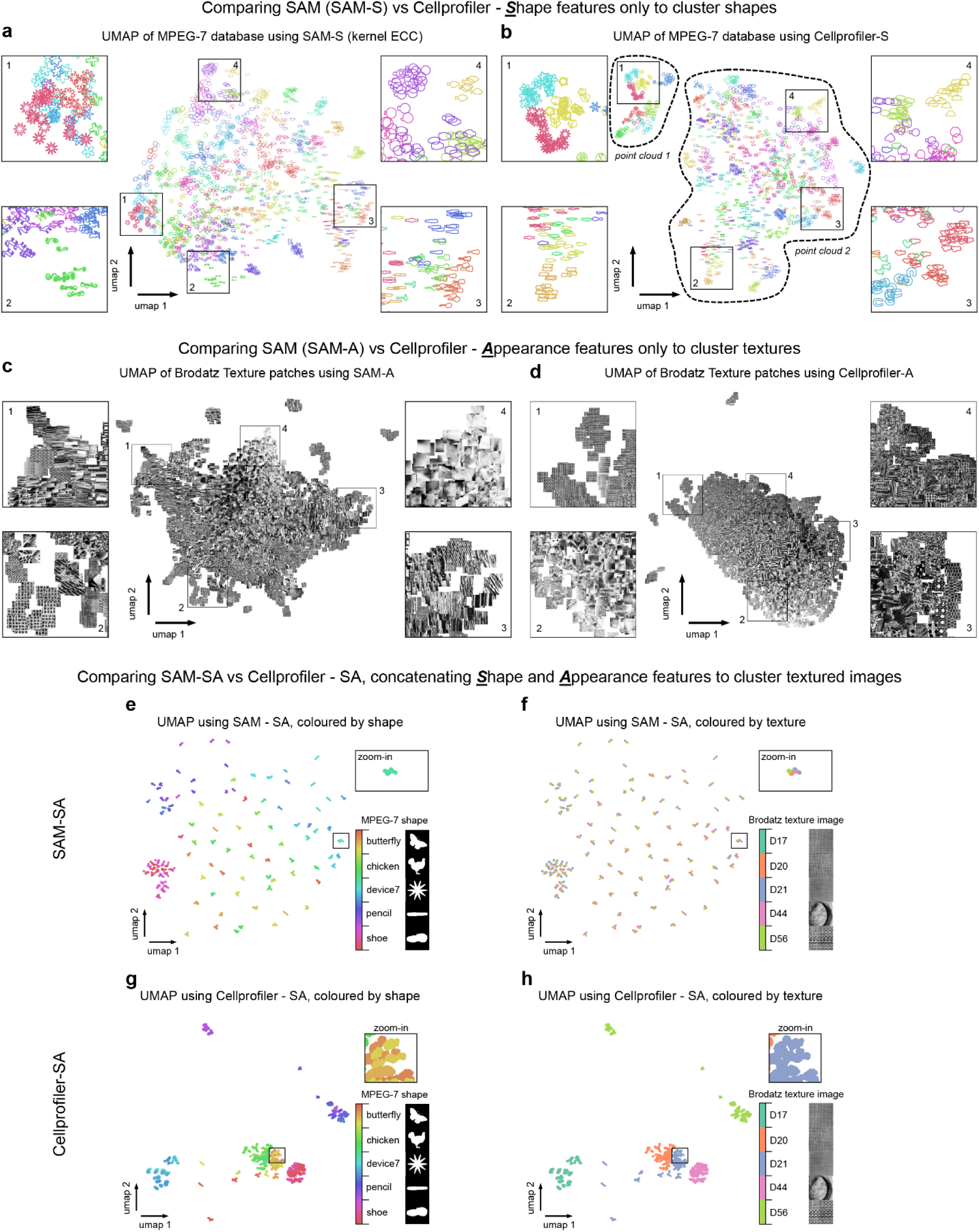
Comparison of using SPOT vs CellProfiler computed shape and appearance features to map phenotypic diversity in computer vision datasets. **a)** Shape phenomic landscape constructed by applying 2D-UMAP to SPOT’s shape-related features, SAM-S (kernel ECC), same as in Fig. 3b. **b)** Shape phenomic landscape constructed by applying 2D-UMAP to CellProfiler’s shape-related features. Each point is an individual shape image in the MPEG-7 dataset colored by its shape category. Similar shapes colocalize to the same local region of the landscape as shown by zoomed-in images 1-4 but are less distinctly separated compared to SAM-S in (a). Moreover, the landscape comprises two point-clouds, indicated by dash black contour lines. **c)** Appearance phenomic landscape constructed by applying 2D-UMAP to SPOT’s appearance-related features, SAM-A, same as in Fig. 3d. **d)** Appearance phenomic landscape constructed by applying 2D-UMAP to CellProfiler-A, CellProfiler’s set of exclusively appearance-related features. Each shape is a 64×64 image patch cropped from the original 512×512 112 Normalized Brodatz texture images. Images cropped from the same original Brodatz texture image and primarily images with the same intensity hue appear to the same local region of the landscape as shown by zoomed-in images 1-4. This contrasts with SAM-A in (c) which organizes primarily by texture. **e,f)** Shape-appearance phenomic landscape constructed by applying 2D-UMAP to the concatenated SAM-S and SAM-A feature sets of the textured shapes created from five selected MPEG-7 shape categories (Fig. 3e) and five randomly selected Normalized Brodatz texture images. Each point is a 128×128 image colored by **e)** its source MPEG-7 shape category and **f)** source Brodatz texture. **g,h)** Shape-appearance phenomic landscape for concatenated CellProfiler-S and CellProfiler-A applied to the same dataset as e,f), colored by **g)** its source MPEG-7 shape category and **g)** source Brodatz texture.

**Supplementary Figure 4a.**
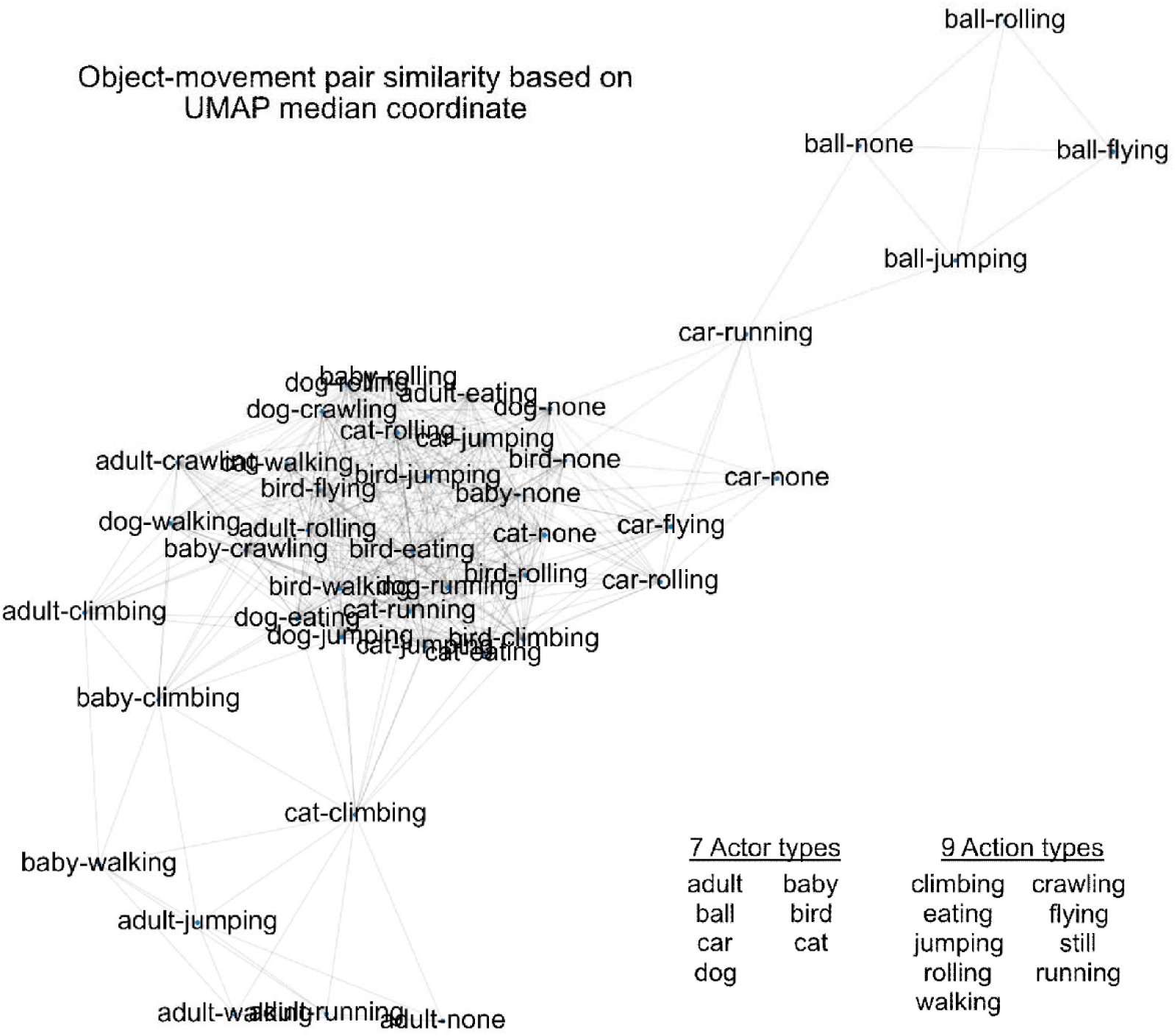
Similarity graph between the 43 unique object-movement classes of the A2D dataset. Graph computed using the pairwise distance between the median 2D-UMAP phenomic landscape coordinates of each unique object-movement class pairing, (Methods). Visualization uses a Kamada-Kawai force-directed layout initialized with the median UMAP coordinates^81^. None refers to the still action type.

**Supplementary Figure 4b.**
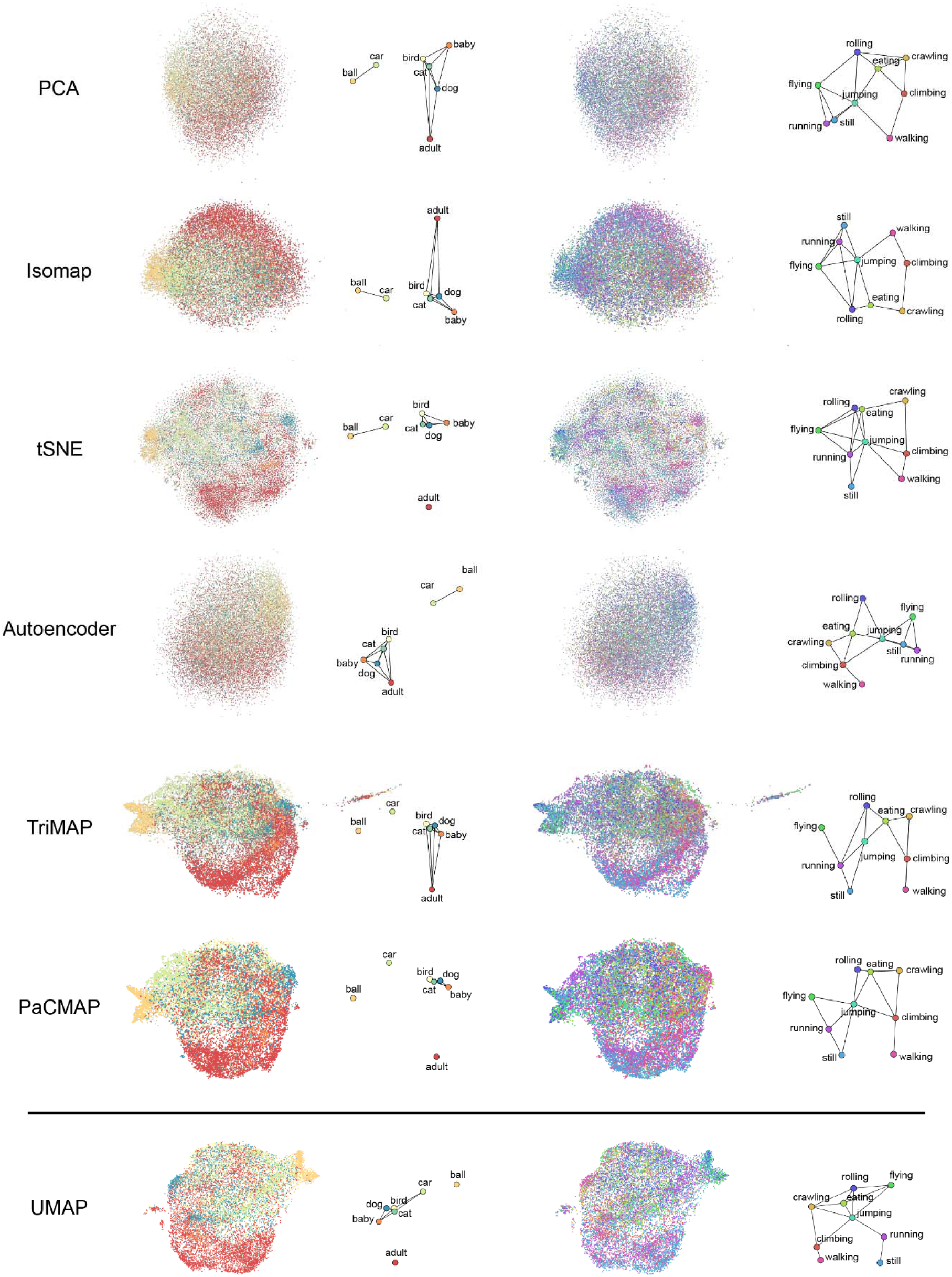
Comparison of different dimensionality reduction techniques and resultant similarity graphs of the seven unique object and nine unique movement classes of the A2D dataset. Computed 2D phenomic landscape with each indicated dimensionality reduction method. Each individual point is an object and is colored by object (first column) and movement (third column) class. Object (second column) and movement (fourth column) similarity graphs were computed using the pairwise distance between the median 2D phenomic landscape coordinates of each dimensionality reduction method (Methods). Graphs were visualized using a Kamada-Kawai force-directed layout initialized with the median UMAP coordinates^81^.

**Supplementary Figure 5.**
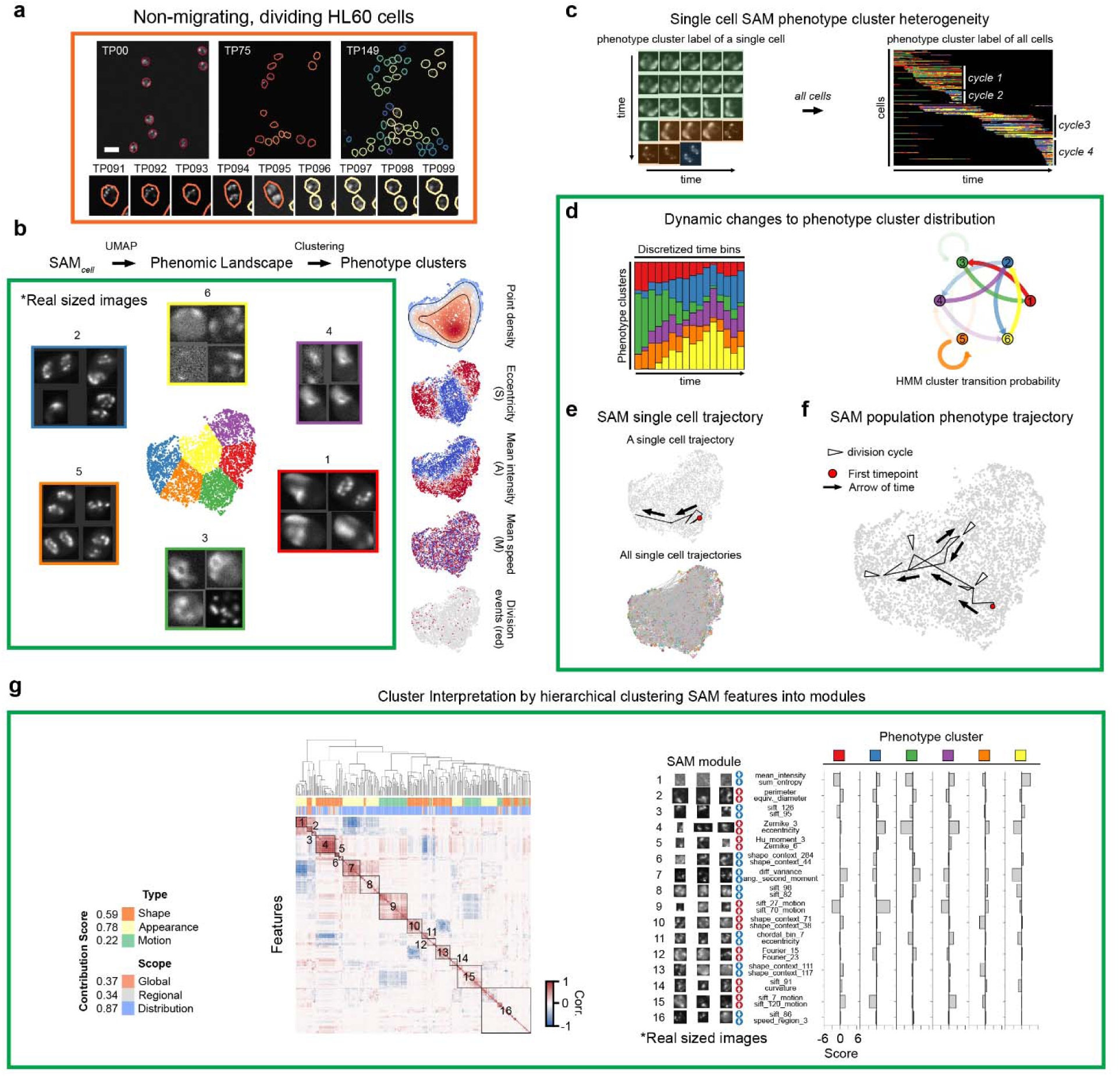
SPOT characterizes phenotypic heterogeneity of single cell division. **a)** Snapshot of the full field-of-view at timepoint 0, 75 and 149 of simulated HL60 cell nuclei with the boundary of each nucleus uniquely colored (top), and snapshots of select cells before or after cell division (bottom). The videos show majority cell division with little migration. Each timepoint is a video frame and is 29 min. Scalebars: 10 µm. **b)** SAM phenomic landscape constructed by applying UMAP to the SAM phenome, followed by identification of phenotype clusters by k-means clustering of 2D-UMAP coordinates using the elbow method. 2×2 image panels show exemplars of the principal phenotypes in each color-coded cluster (left, Methods). Local point density of mapped cell instances, whereby each point, representing a cell instance, is colored to indicate low-to-high (blue-to-red) measured values in global SAM features of shape (eccentricity), appearance (intensity), and motion (speed), (right, first to fourth panel, top-to-bottom). Instances were also colored discretely as red or grey to indicate if it was the first timepoint after cell division (right, fifth panel). **c)** Mapping a single cell tracked over time (left) and all continuously tracked cells (right) into the SAM phenomic landscape of **b)** and coloring each temporal instance by the corresponding phenotype cluster. **d)** Stacked barplot showing the relative frequency of each phenotype cluster over discrete time bins (left). Graph showing the Hidden Markov Model (HMM) inferred transition probability (Methods) of a cell transitioning to another phenotype cluster in the next timepoint, given its phenotype cluster label in the current timepoint (right). Arrows colored by the source cluster. The more transparent the arrow the smaller the probability of transition. **e)** Single SAM phenotype trajectory summarizing the temporal phenotype dynamics of a single HL60 cell (top) and the SAM phenotype trajectory of all cells (bottom) with starting timepoint colored red. Black arrow shows the directionality of time. **f)** Single population-level SAM phenotype trajectory summarizing the temporal evolution and phenotypic diversity across all cells. Starting timepoint is colored red. Black arrows show the directionality of time. White arrows depict a cell division cycle. **g)** Contribution score of shape, appearance and motion, and global, regional and distributional features in explaining the dataset variance defined as the absolute value of the first principal component (left). Automated hierarchical clustering of the covariation between SAM features to identify principal SAM feature modules (middle, modules outlined and numbered along the diagonal). Expression of each SAM module (labeled ‘score’ in barplot) in each phenotype cluster (right). Each module is depicted with their top three most representative images, its top driving SAM features and whether each feature is enriched (up arrow) or depleted (down arrow).

**Supplementary Figure 6.**
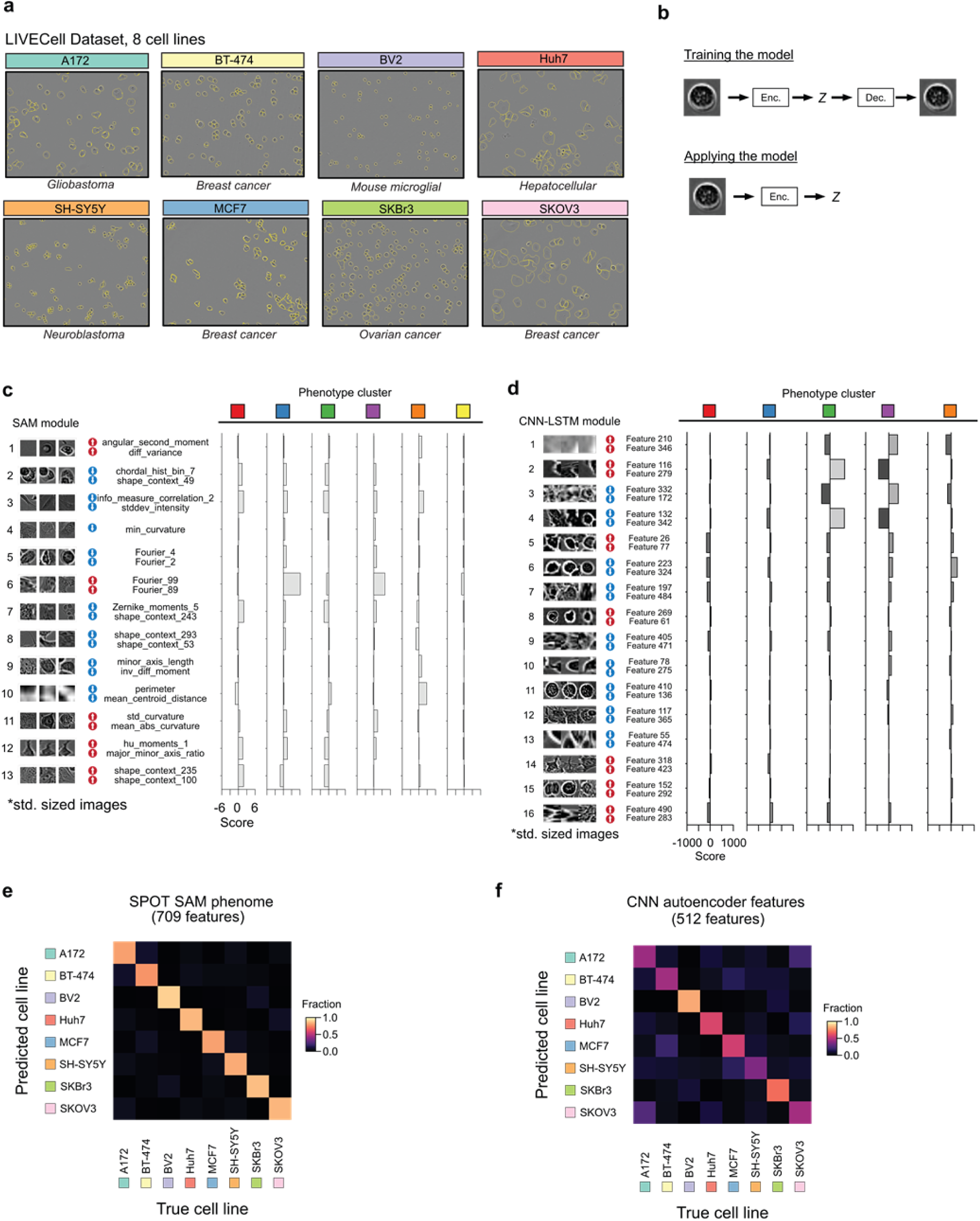
SPOT analysis of LIVECell dataset using SAM and CNN autoencoder features. **a)** Example phase-contrast images, with provided manual cell outlines (yellow) for each of the 8 cell lines in the LIVECell dataset. **b)** Schematic of training a convolutional neural network autoencoder to extract features from image crops (resized to standard size, 64 x 64 pixels same as that used in SPOT analysis) using reconstruction (top), and then applying the trained encoder to extract features (bottom). **c,d)** Expression of each module (labeled as ‘score’ in barplots, Methods) in each phenotype cluster following hierarchical clustering of **c)** SAM and **d)** CNN features into modules. Each module is depicted with its three selected most representative images, its top driving features and whether the feature is enriched (up arrow) or depleted (down arrow). **e)** Confusion matrix using the SPOT filtered SAM phenome after preprocessing (parameters set by the train split) and feature normalization on the test split. **f)** Confusion matrix using the CNN encoder generated features as-is on the same test split as the SAM phenome, with no feature selection or normalization.

**Supplementary Figure 7.**
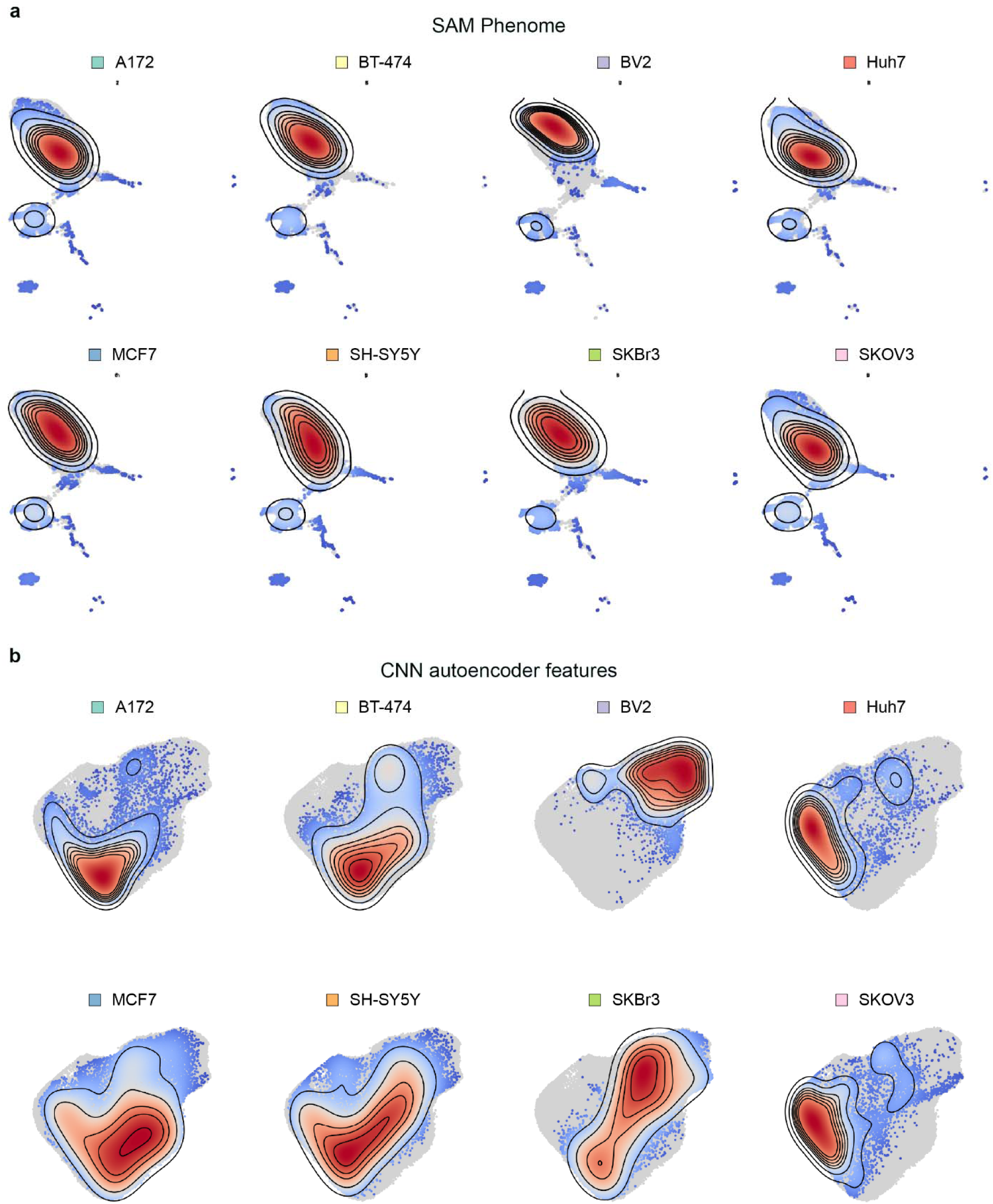
Density heatmap of individual cell lines when mapped into the SAM phenome or CNN features phenomic landscape. **a)** Heatmap of the local point density of cells mapped into the phenomic landscape generated by applying UMAP to the SAM phenome. **b)** Same as a) after applying UMAP to the CNN features. Contour lines depict mean + 1, 2, 3, 3.5, 4, 4.5 and 5 standard deviations of the point density.

**Supplementary Figure 8.**
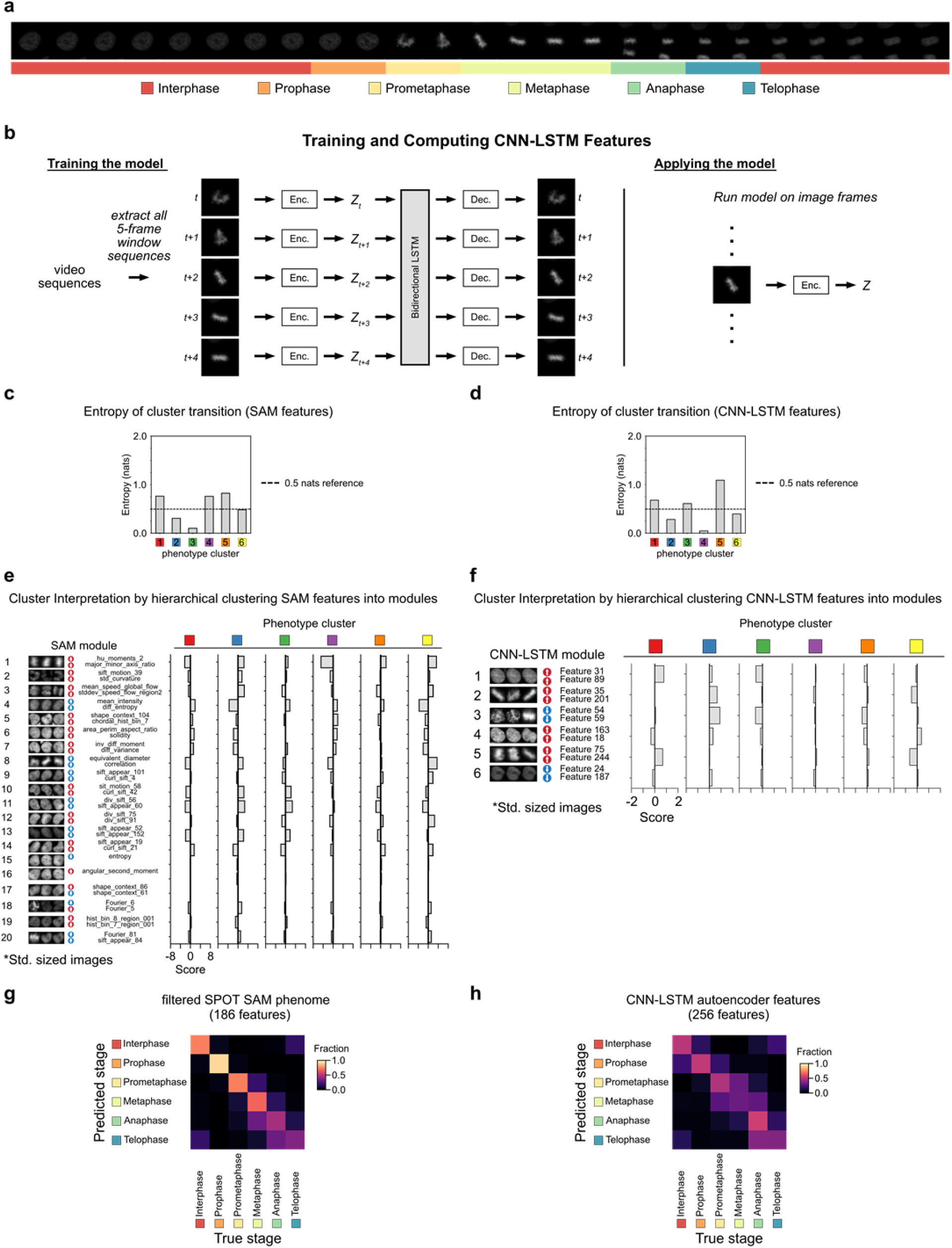
SPOT analysis of HeLa cell histone H2B dataset using SAM and CNN-LSTM autoencoder features. **a)** Representative video from the HeLa cell histone H2B dataset for a single cell, showing the changes in nuclear morphology as the cell divides and the corresponding manually annotated staging, depicted as a colorbar below. A unique color is ascribed to each of the six stages. **b)** Schematic of training the Convolutional Neural Network – Long Short-Term Memory (CNN-LSTM) autoencoder to learn to extract temporally-informed features trained on 5-frame image crop sequences (each crop resized to standard size, 64 x 64 pixels same as that used in SPOT analysis) by reconstruction (left), and then applying the trained CNN encoder to extract features from individual single timepoint image crops (right). **c), d)** Shannon entropy of cluster transition for each phenotype cluster found by SPOT analysis of the SAM phenome or CNN-LSTM features respectively. **e)** Automated hierarchical clustering of the covariation between SAM features to identify principal SAM feature modules. The expression of each SAM module (labeled as ‘score’ in barplots, Methods) in each phenotype cluster. Each module is depicted with its three selected most representative images, its top driving SAM features and whether the feature is enriched (up arrow) or depleted (down arrow) (see Methods). **f)** Automated hierarchical clusters of the covariation between CNN-LTSM features to identify principal feature modules, as for **e). g)** Confusion matrix using the SPOT filtered SAM phenome after preprocessing (parameters set by the train split) and feature normalization on the test split. **h)** Confusion matrix using the trained CNN-LSTM encoder generated features as-is on the same test split as the SAM phenome, with no feature selection or normalization.

**Supplementary Figure 9.**
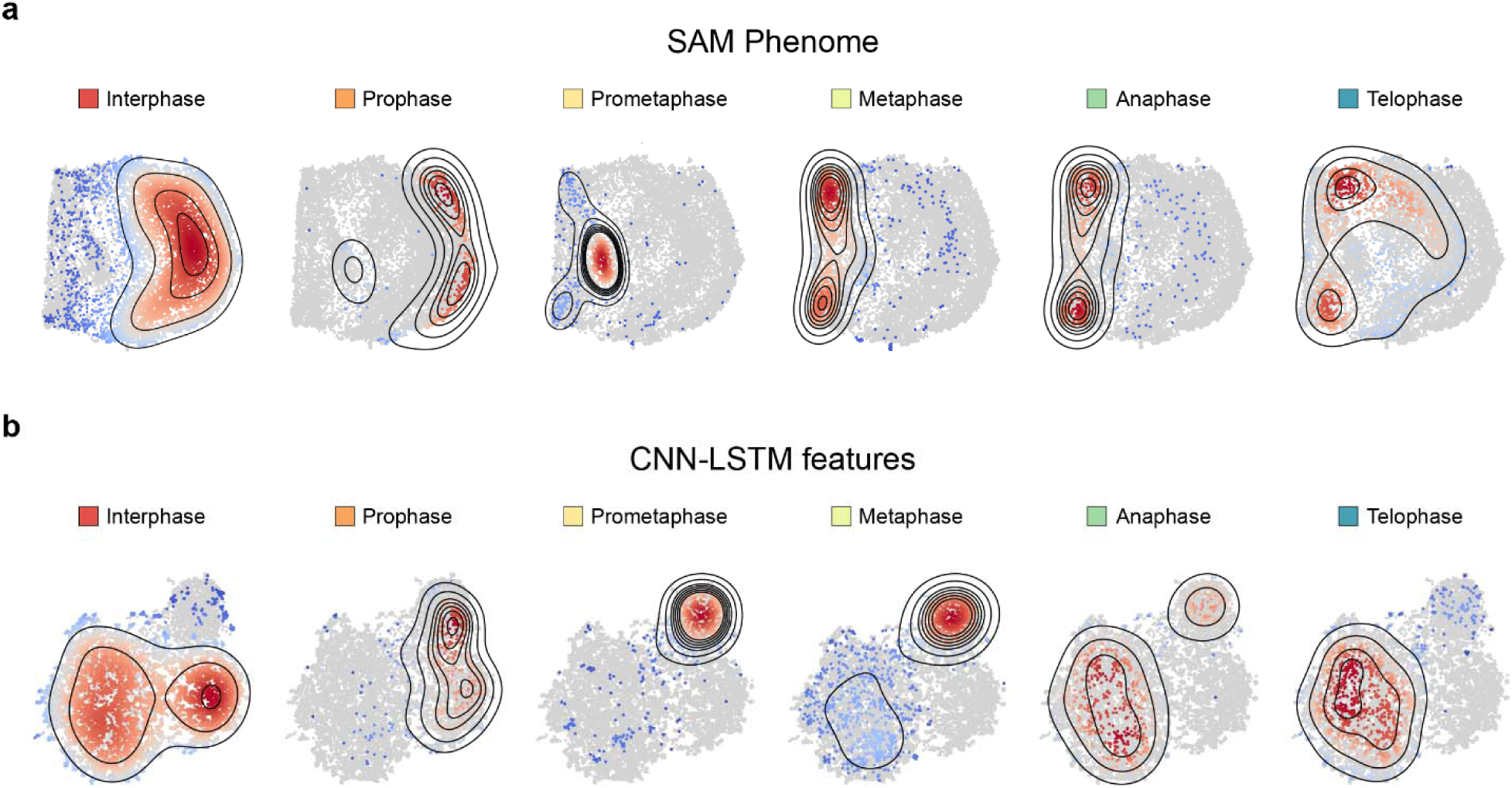
Density heatmap of cell division stage when mapped into the SAM phenome or CNN-LSTM features phenomic landscape. **a)** Heatmap of the local point density of cells mapped into the phenomic landscape generated by applying UMAP to the SAM phenome. **b)** Same as a) after applying UMAP to the CNN-LSTM features. Contour lines depict mean + 1, 2, 3, 3.5, 4, 4.5 and 5 standard deviations of the point density.

## SUPPLEMENTAL TABLES

**Supplementary Table 1.** Summary of the 2185 computed Shape, Appearance and Motion (SAM) features which constitute the SAM phenome in this study.

**Supplementary Table 2.** Performance metrics of SPOT’s SAM features and CellProfiler’s features on computer vision datasets.

**Supplementary Table 3.** Classifer performance for predicting cell line in the LIVECell dataset using SPOT’s SAM phenome and CNN autoencoder features learnt directly from images.

**Supplementary Table 4.** Classifer performance for predicting cell division stages using SPOT’s SAM phenome and CNN-LSTM autoencoder features learnt directly from video sequences for a timelapse image dataset of HeLa cell histone H2B.

## SUPPLEMENTAL MOVIES

**Supplementary Movie 1.** Reference cell segmentation and tracking of migrating glioblastoma-astrocytoma U373 and dividing HL60 leukaemia cells from the 2D cell tracking challenge. Videos show the curated reference individual cell segmentations and tracks for both videos in the training data of U373 and HL60 cells used for SPOT analysis. Individually tracked cells are colored uniquely. Where cells move partially out of the field of view, these individual instances are removed prior to analysis and are not labeled. Consequently, if a cell leaves the field of view and later returns, the separate tracks are treated as separate entities for Hidden Markov Model (HMM) analysis. Movie frames are acquired every 15 min for U373 and every 29 min for HL60. Scalebar: 50 μm for U373, 10 μm for HL60.

